# CD45 pre-exclusion from the tips of microvilli establishes a phosphatase-free zone for early TCR triggering

**DOI:** 10.1101/2020.05.21.109074

**Authors:** Yunmin Jung, Lai Wen, Amnon Altman, Klaus Ley

**Affiliations:** Center for Autoimmunity and Inflammation, La Jolla Institute for Immunology, La Jolla, CA, USA; Center for Cancer Immunotherapy, La Jolla Institute for Immunology, La Jolla, CA, USA; Department of Bioengineering, University of California San Diego, La Jolla, CA, USA

**Keywords:** CD45, TCR signaling, Pre-exclusion, Expansion microscopy, Microvilli, Plasma membrane, Cholesterol, Lipid rafts, Calcium influx, Immunological synapse

## Abstract

The tyrosine phosphatase CD45 is a major gatekeeper for restraining T cell activation. Its exclusion from the immunological synapse (IS) is crucial for TCR signal transduction. Here, we used expansion super-resolution microscopy to reveal that CD45 is pre-excluded from the tips of microvilli on primary T cells prior to antigen encounter. This pre-exclusion was diminished by depleting cholesterol or by engineering the transmembrane domain of CD45 to increase its membrane integration length, but was independent of the CD45 extracellular domain. We further show that brief microvilli-mediated contacts can induce Ca^2+^ influx in mouse antigen-specific T cells engaged by antigen-pulsed APCs. We propose that the absence of CD45 phosphatase activity at the tips of microvilli enables or facilitates TCR triggering from brief T cell-APC contacts before formation of a stable IS, and that these microvilli-mediated contacts represent the earliest step in the initiation of a T cell adaptive immune response.

**Graphical abstract:** 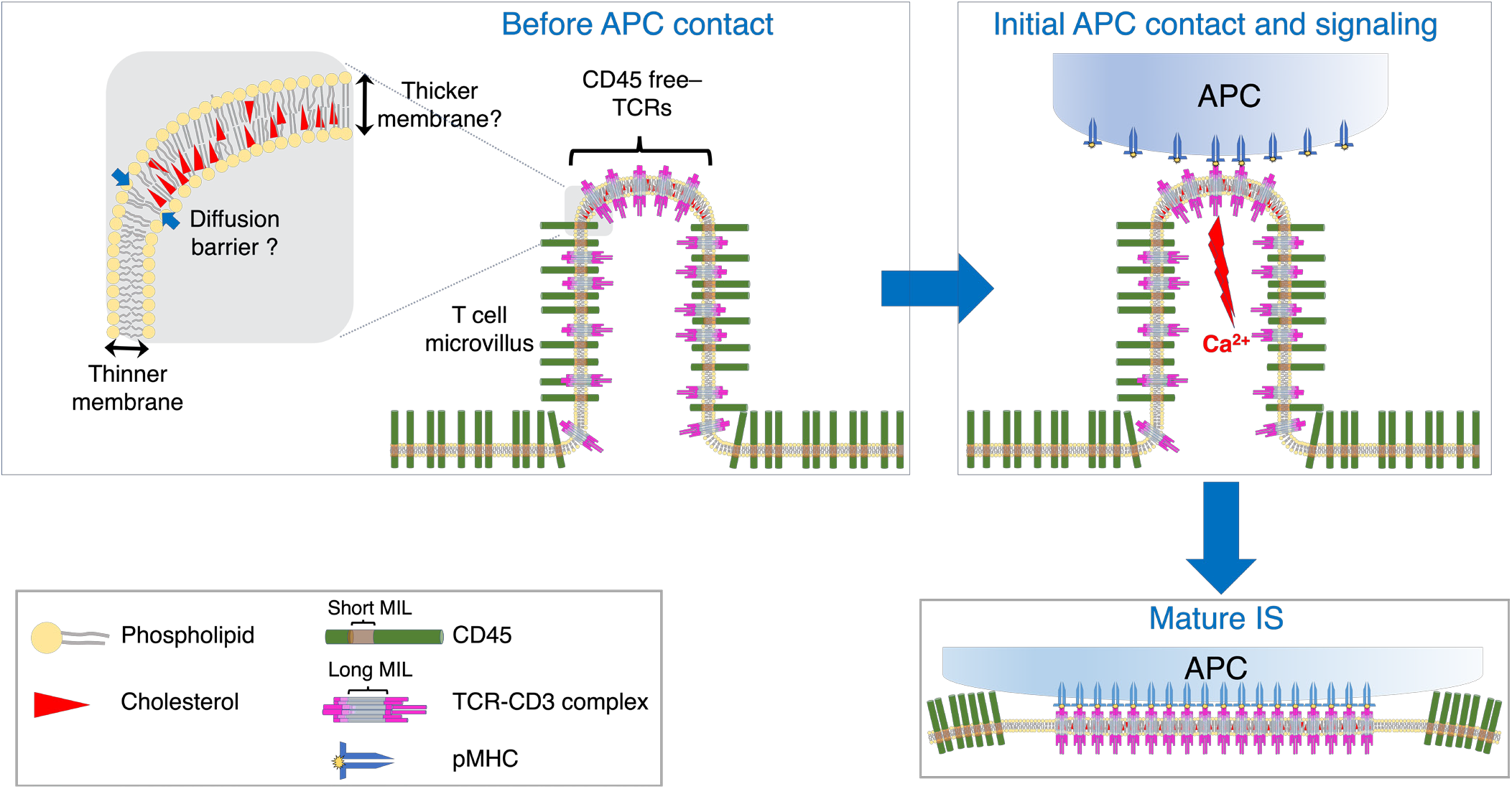

## Introduction

Transmembrane tyrosine phosphatase CD45 is expressed on all nucleated hematopoietic cells and is one of the most abundantly expressed membrane proteins on lymphocytes (Penninger et al., 2001; Tonks et al., 1988). While its cytoplasmic domain, including the catalytic domain, shares 95% homology among mammalian species, the highly glycosylated extracellular domain, which includes several splicing variants, shares only 35% homology (Thomas, 1989). CD45 is known to be critical for restraining the adaptive T cell response by negatively regulating T cell receptor (TCR) signaling, unless the T cell encounters a high-affinity cognate antigen (Courtney et al., 2019; Sibener et al., 2018).

The molecular mechanism of TCR triggering in the immunological synapse (IS) has been explained by size-dependent kinetic segregation (Shaw and Dustin, 1997). Once TCRs binds to its cognate antigens presented by major histocompatibility complex (MHC) molecules on an antigen presenting cell (APC), CD45 molecules are excluded from the tight contact region, shifting the equilibrium to the phosphorylated state of the immunoreceptor tyrosine-based activation motifs (ITAMs) in the TCR-CD3 complex and allowing productive T cell activation. CD45 has a long extended extracellular domain, which exceeds the length of the TCR-pMHC complex (Penninger et al., 2001). Therefore, the long extracellular domain of CD45 that cannot be accommodated within the tight contact interface between T cells and APCs is segregated from the interface (Schmid et al., 2016). TCR engagement by its cognate MHC-bound peptide is sufficient to drive CD45 exclusion in the absence of any downstream signaling (James and Vale, 2012). Recent studies using super-resolution optical techniques have revealed that CD45 exclusion is initiated from microvilli-mediated close contacts that expand as the IS enlarges and matures (Chang et al., 2016; Razvag et al., 2018). CD45 exclusion is crucial for initial TCR triggering events (Chang et al., 2016; Razvag et al., 2018; Varma et al., 2006) and it correlates with effective TCR signaling (Sibener et al., 2018; Varma et al., 2006). Artificially induced CD45 exclusion can initiate TCR triggering in a non-immune cell reconstitution system (James and Vale, 2012). CD45 exclusion also plays a key role in B cell receptor (BCR) signaling and high-affinity immunoglobulin E receptor (FcεRI) signaling in mast cells and basophils (Coughlin et al., 2015; Depoil et al., 2008; Felce et al., 2018).

Microvilli are thin actin-based membranous projections that are found on all immune cells (Majstoravich et al., 2004). Several recent studies have highlighted the role of microvilli in T cell activation. TCRs are highly enriched in the microvilli of both resting and effector primary T cells, suggesting that microvilli act as effective sensors for antigenic moieties (Jung et al., 2016). Lattice light-sheet microscopy revealed that dynamic motions of microvilli enable mouse T cells to scan the majority of the opposing surfaces of an APC within one minute (Cai et al., 2017). Quantitative Ca^2+^ imaging combined with interference reflection microscopy (IRM) showed that an intracellular Ca^2+^ response was triggered as early as one minute after initial contact between a T cell and an activating antibody-coated glass surface, preceding actin remodeling and spreading (Fritzsche et al., 2017). Initial microvilli-mediated isolated contacts and local CD45 exclusion from those areas were sufficient for robust TCR signaling (Razvag et al., 2019). Thus, microvilli-mediated initial contacts likely represent an early crucial step in the decision making process of T cells whether to respond to an antigen.

The plasma membrane (PM) contains diverse types of phospholipids and other lipids such as sterols and sphingolipids, which account for variation in membrane thickness and composition (Harayama and Riezman, 2018; Mitra et al., 2004; Niemela et al., 2007). Cholesterol is known to accumulate in the inner membrane leaflet at negative curvatures (Cebecauer et al., 2018; Garcia-Arribas et al., 2016), and this high cholesterol content causes an increase in the thickness of the lipid bilayer (Favela-Rosales et al., 2019; Niemela et al., 2007; Smondyrev and Berkowitz, 1999). The thickness of pure phospholipid bilayers is ~28-35 Å depending on the types of phospholipid (Harroun et al., 1999; Hills and McGlinchey, 2016). However, the cell membrane at lipid raft domains, which are enriched in sphingomyelin and cholesterol is 4~8 Å thicker than that of phospholipid bilayers lacking these lipids (Favela-Rosales et al., 2019; Niemela et al., 2007; Smondyrev and Berkowitz, 1999). Lipid partitioning in the PM can determine, in turn, the sorting of membrane proteins (Sengupta et al., 2019). It has been reported that approximately 95% of CD45 is found in the non-raft fraction of cell membrane (Edmonds and Ostergaard, 2002).

Here, we set out to examine whether the TCR and CD45 segregate from each other in T cell microvilli and, furthermore, whether CD45 (but not the TCR) may be pre-excluded from the tips of microvilli prior to T cell engagement by APCs, potentially as a result of diffusion barriers formed by cholesterol accumulation and, hence, thicker membrane at the tip region. Using advanced microscopy, namely, 4x expansion microscopy (Chen et al., 2015; Tillberg et al., 2016) followed by Airyscan laser scanning confocal imaging (Huff, 2015), which yielded an effective resolution of 30-40 nm, we analyzed CD45 localization within the tips *vs*. the columns of microvilli. We report that CD45 is pre-excluded from the tips of microvilli in T cells prior to their early search and scanning of APCs (Cai et al., 2017), and confirmed this using an independent approach (STORM super-resolution microscopy). CD45 pre-exclusion was found in different types of lymphocytes, including CD4^+^ and CD8^+^ T cells, regulatory T cells (Tregs), and B cells. Furthermore, microvilli-mediated engagement of APCs by antigen-specific T cells during the early scanning stage (Cai et al., 2017) resulted in rapid Ca^2+^ influx. Thus, CD45 pre-exclusion from the tips of microvilli, which exists prior to any APC contact effectively establishes a phosphatase-free zone at the tips of lymphocyte microvilli, poising them for an immediate response upon ligand engagement.

## Results

### CD45 is pre-excluded from the tips of microvilli in human and mouse lymphocytes

Under conventional microscopy, the thin and short protrusive structures of microvilli appear as blurred short spikes on the cell surface. Therefore, mapping the distribution of molecules of interest in relation to microvilli and their sub-regions requires super-resolution imaging techniques. Although Airyscan imaging offers ~1.7-fold better resolution than that achieved by conventional light microscopy, Airyscan was not sufficient to resolve the localization of labeled CD45 on the microvilli (Figure 1A). To achieve better resolution, we applied 4x expansion microscopy (Chen et al., 2015; Tillberg et al., 2016) combined with Airyscan confocal imaging (4xExM-Airyscan). Since the resolving power of expansion microscopy is linearly increased as a function of the expansion factor of the swellable hydrogel, a combination of these two techniques can achieve a ~1.7*4=6.8-fold higher resolution than the diffraction limit. This approach allowed imaging of the entire cell, making it possible to obtain cross sectional images of hundreds of microvilli, thus enabling us to unravel the nanoscale molecular distribution of CD45. We find that CD45 is excluded from the tips of microvilli on freshly isolated, unstimulated human CD4^+^ T cells (Figure 1B). Post-staining with the membrane staining dye, FM 4-64FX, confirmed that CD45 was excluded from the tips of microvilli (Figure 1C). As a control, we imaged the localization of L-selectin, which is known to be expressed on T cell microvilli (Jung et al., 2016; von Andrian et al., 1995), and found that, unlike CD45, which was largely excluded from the tips, L-selectin was highly concentrated on the tips. (Figures 1D-F). The CD45 exclusion did not correlate with the expression level of L-selectin (Figure S1). We confirmed CD45 pre-exclusion from the microvilli tips by using another super-resolution imaging technique, *i.e*., stochastic optical reconstruction microscopy (STORM) (Figure S2). Importantly, this CD45 pre-exclusion from the tips of microvilli in unstimulated T cells is distinct from the previously reported ligand-induced exclusion (Razvag et al., 2018). We observed similar CD45 pre-exclusion in human CD8^+^ T cells, (Figure S3A) and mouse CD4^+^ T cells (Figure S3B), as well as in regulatory T (Treg) cells sorted from *Foxp3*^YFP-Cre^ transgenic mice (Figure S3C). Similar CD45 exclusion was also found in both human and mouse B cells (Figures S3D and S3E). These findings indicate that CD45 pre-exclusion from the tips of microvilli is a universal phenomenon in lymphocytes.

**Figure 1.**
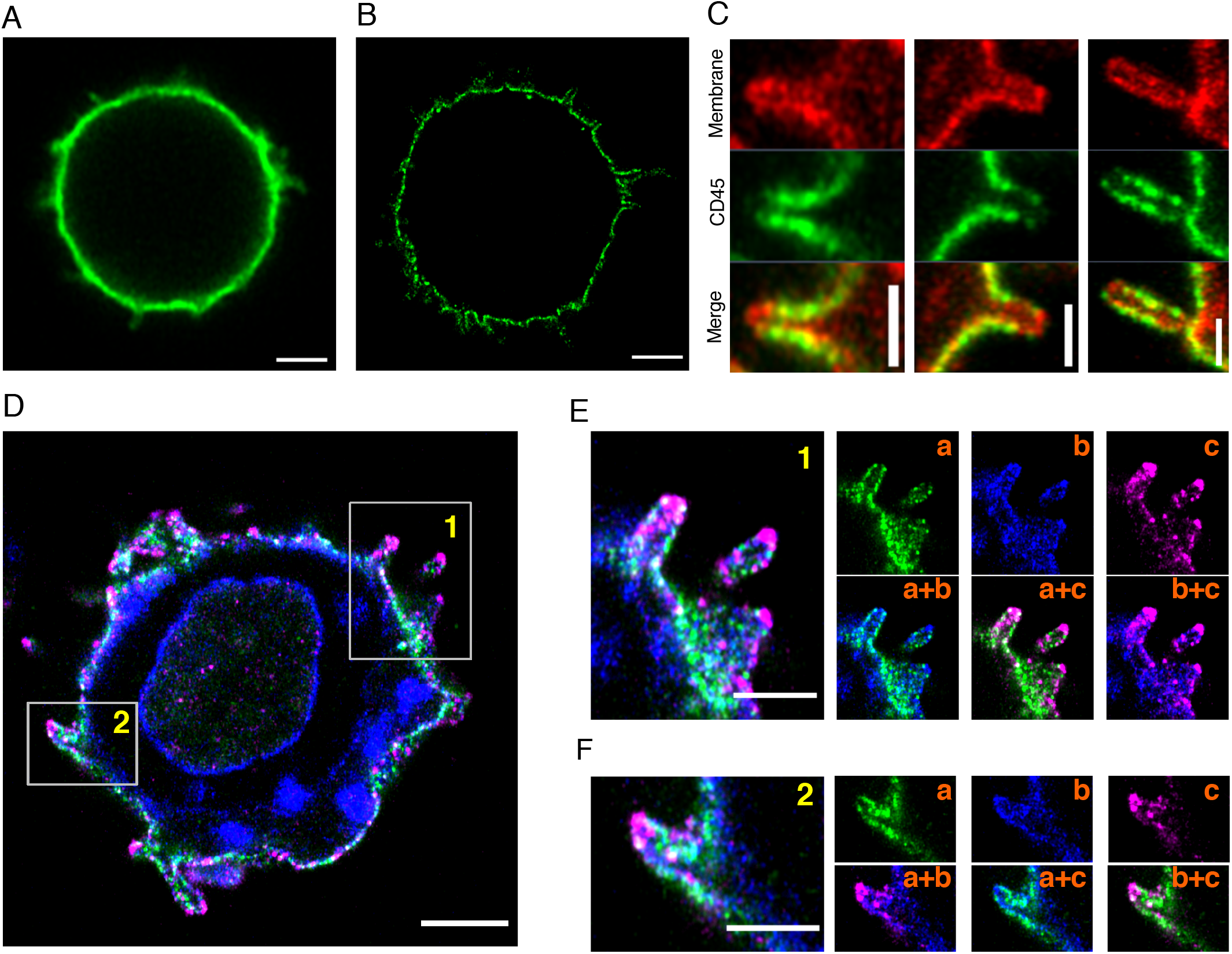
Pre-exclusion of CD45 at the tips of microvilli on human CD4^+^ T cells. (A) A representative Airyscan image of a human CD4^+^ T cell labeled with Alexa Fluor 488 (AF488) conjugated anti-CD45 antibody. (B) A representative 4x-ExM-Airyscan image of a similarly labeled cell. (C) Representative 4x-ExM-Airyscan images of microvilli on human CD4^+^ T cells stained with anti-CD45-AF488 (green) and FM 4-64FX membrane dye (red). (D) A representative 4x-ExM-Airyscan image of a human CD4^+^ T cell stained with anti-CD45-AF488 (green), FM 4-64FX membrane dye (blue) and anti-L-selectin-AF568 (magenta). (E-F) Magnified images of areas 1 and 2 marked by the rectangles in (D). Individual channel images of CD45 (a), membrane (b), and L-selectin (c) (right, upper panels) and merged images of (a) and (b), (a) and (c) and (b) and (c) (right, lower panels). These images are representative of 40 analyzed cells. Scale bars in A, B, D: 2 μm; in C: 500 nm; E, F: 1 μm. The scale bars are corrected by the expansion factor except for (A).

### CD45-free TCRs and BCRs are present on the tips of microvilli

TCRs are enriched on microvilli of quiescent human and mouse T cells (Jung et al., 2016; Cai et al., 2017). Combining these findings with our observation of CD45 pre-exclusion, CD45-free TCRs are expected to be enriched at the tips of microvilli. As shown in Figures 2A-C, CD3 molecules were highly enriched on the entire microvilli surface compared to the cell body, while CD45 molecules were excluded from the tips. We quantified the relative intensities of these two molecules by determining the ratiometric generalized polarization (GP) (Table S1) (Yu et al., 1996). A GP value of −1 represents 100% exclusion of CD45, and +1 represents 100% CD45 enrichment. Figure 2D shows the segmented areas for outside-cell background (OutBG), inside-cell background (InBG), cell-body area (CB), microvilli area (MV), and microvilli tips of the representative cell shown in Figures 2A-C. Figure 2E shows color-coded GP values where yellow is −1 and blue is +1. The distributions of GP values within the MV (Figure 2F) and CB (Figure 2G) areas of the cell shown in Figure 2 revealed that negative GP values indicative of CD45 exclusion were mostly found within the MV area rather than in the CB area.

**Figure 2.**
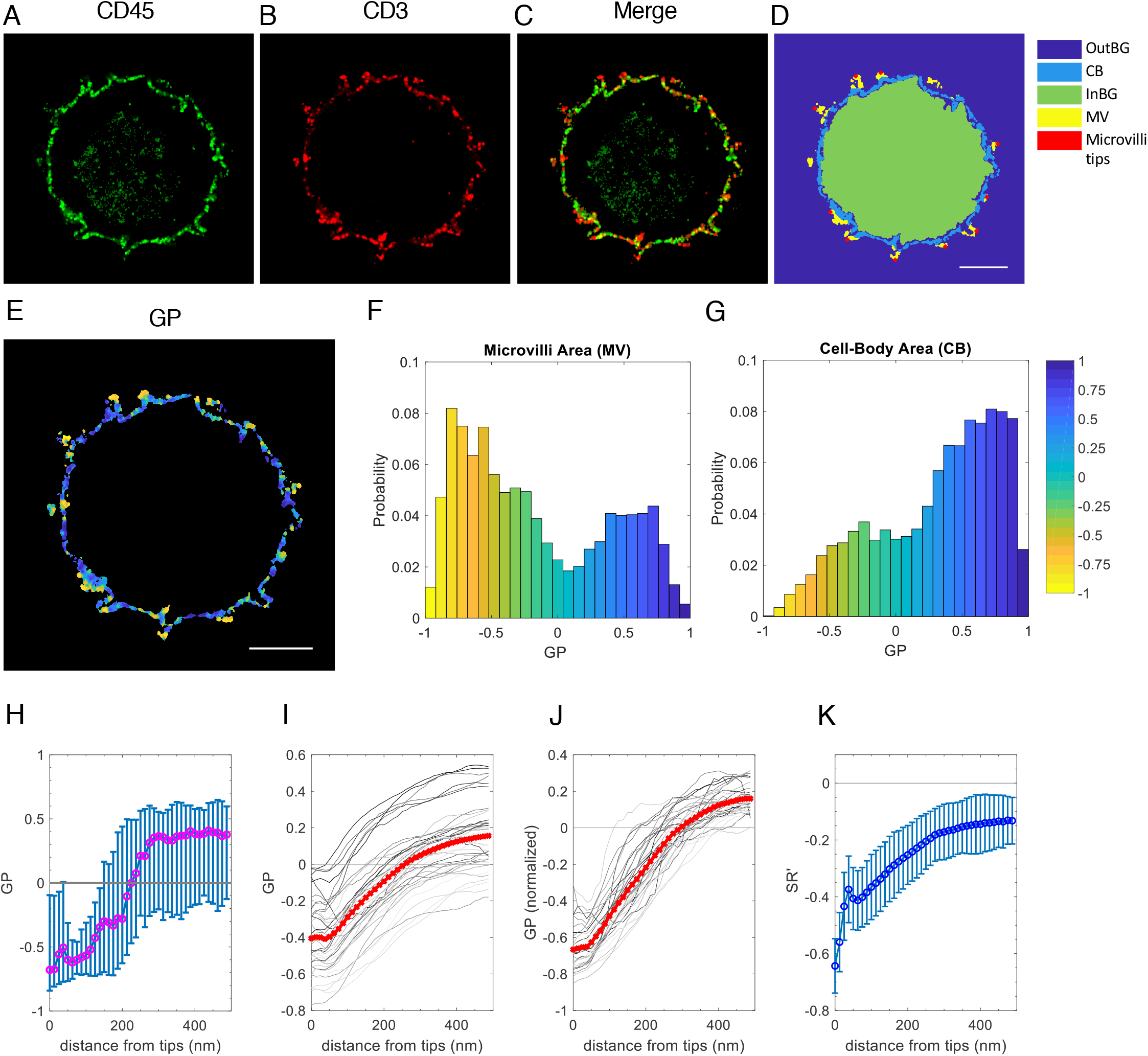
Presence of CD45-free CD3 molecules on the tips of microvilli in human CD4 T^+^ cells. (A-C) Representative 4x-ExM-Airyscan images of a CD4^+^ T cell labeled with anti-CD45-AF488 (green; A), anti-CD3-CF633 (red; B) and the merged image (C). (D) Area map of the cell in (A-C) segmented for outside-cell background (OutBG), cell body (CB), inside-cell background (InBG), microvilli (MV) and positions of the individual microvilli tips (microvilli tips). (E) The GP image between A and B. Scale bars in (D, E): 2 μm. (F, G) Normalized distribution of the GP values within the MV (F) and CB (G) areas of the cell shown in (D). (H) Median GP values of image (E) segmented by distance from the location of microvilli tips shown in (D) were plotted as a function of the distance from the microvilli tips. Error bars represent the 25th and 75th percentiles of the data. (I) The median GP values of 35 cells were plotted as a function of the distance from microvilli tips. Each line represents data collected from 20 z-plane images of a cell; the red line is the mean of the 35 plots. (J) Normalized GP values for each line in (I). (K) The mean *SR’* values between the CD45 and CD3 images segmented by distance obtained from 35 cells were plotted as a function of the distance from microvilli tips. Error bars represent the standard deviation (SD).

Next, we correlated the spatial distribution of GP values to the position of microvilli tips. GP values for each pixel were grouped by distance from tips with a bin size of 12.5 nm, followed by plotting the median GP values of each group as a function of the distance (Figure 2H), where 0 is the position of tips. GP values calculated between the CD45 and CD3 were most negative at the microvilli tips (Figure 2E). Figure 2I shows the individual median GP plots of 35 cells as a function of distance from the microvilli tips. Although the expression levels of molecules varied among cells, the most negative GP values were found near the microvilli tips. This result held up after normalization of mean GP values to zero for all cells (Figure 2J).

To test whether CD45 exclusion was robustly detected independent of the intensity variation, we applied a second, new method that we call segment-correlation coefficient, *SR’*, between the signals from two channels (Table S1 and Figure S4). SR’ quantifies the contribution of negative or positive image correlation in a segmented area regardless of variations in signal intensities among cells. *SR’* values from simulated data for validation (Figures S4A-F) and from the segmented data by distance or GP values of a representative cell showed that *SR’* faithfully displays a correlation between two channels within the segmented area, and it is sensitive to the GP value between the channels. In Figure 2K, the mean *SR’* as a function of distance from the microvilli tips of 35 cells segmented by distance from the microvilli tips showed that the *SR’* values were significantly negative when the distances were less than 50 nm, which matches the radius of microvilli (Jung et al., 2016; Majstoravich et al., 2004). Since the majority of GP values near the tips were mostly negative (Figures 2E-J), this anti-correlation between CD45 and CD3 clearly indicates that CD45-free TCRs were highly enriched at the tips of microvilli.

Next, we investigated the distribution of the B cell receptor (BCR) *vs*. CD45 in B cell microvilli. A previous study showed that ICAM1 and MHC-II are expressed on microvilli while LFA1 and CD40 are excluded (Greicius et al., 2004); however, BCR localization in microvilli has not been reported. We found that CF633-labeled BCR (IgD and IgM) molecules on the cell surface were highly enriched on microvilli (Figure 3B and S5B). Similar to TCRs, CD45-free BCR molecules were highly enriched at the tips of microvilli, where CD45 molecules were largely excluded (Figures 3 and S5). These results indicate that TCRs and BCRs at the tips of microvilli are mostly devoid of CD45 molecules.

**Figure 3.**
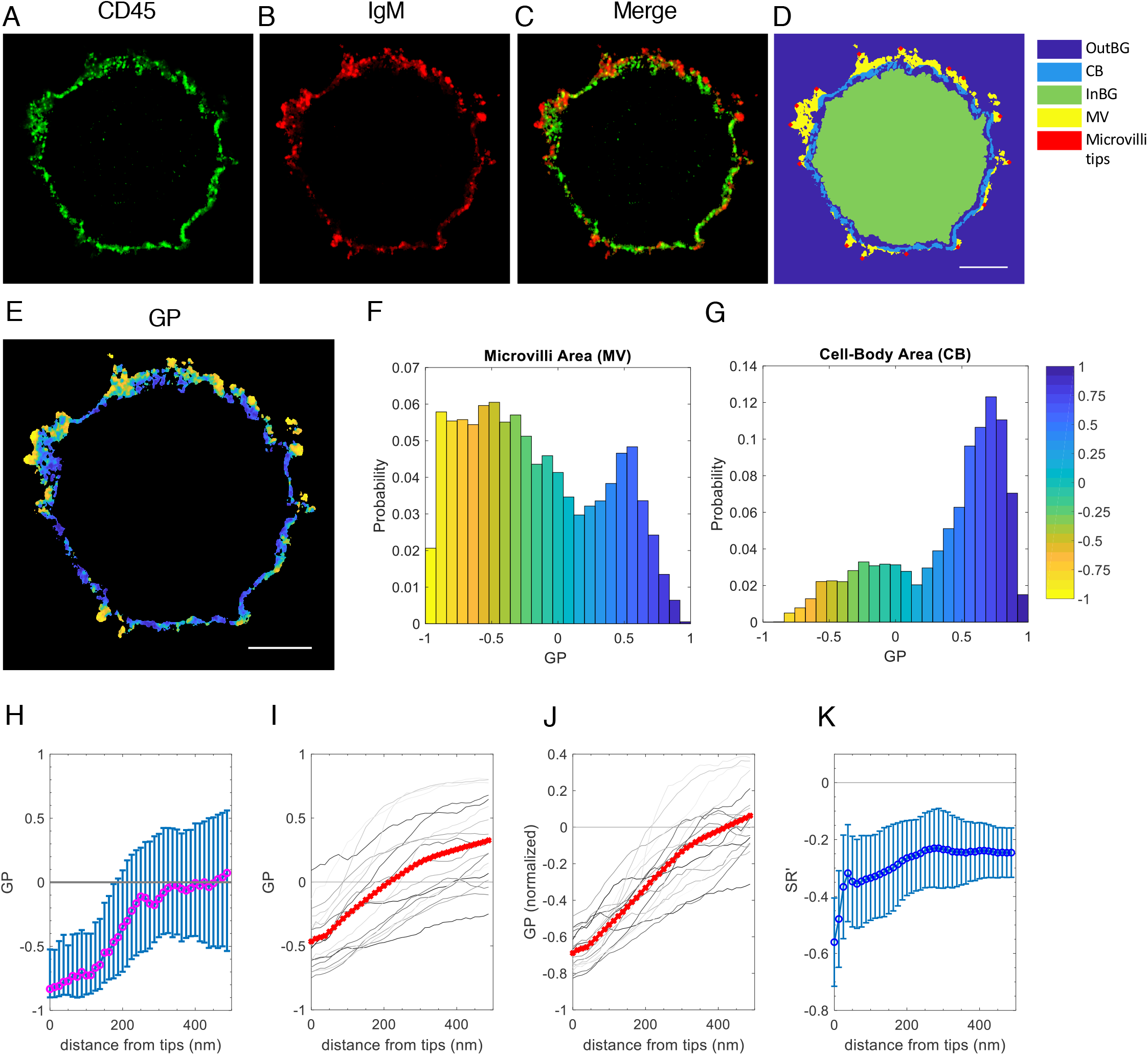
CD45-free membrane-bound IgM molecules on the tips of microvilli of human B cells. (A-C) A representative 4x-ExM-Airyscan image of a B cell labeled with anti-CD45-AF488 (green; A), anti-IgM-CF633 (red; B), and the merged image (C). (D) The area map segmented for OutBG, CB, InBG, MV, and microvilli tips of the images in (A-C). (E) The GP image between A and B. Scale bars in (D, E): 2 μm. (F, G) Normalized distribution of the GP values within the MV (F) and CB (G) areas of the cell shown in (D). (H) Median GP values of the image (E) segmented by distance from the location of microvilli tips shown in (D) were plotted as a function of the distance from the microvilli tips. Error bars represent the 25th percentile and 75th percentile of the data. (I) The median GP values of 19 cells were plotted as a function of the distance from microvilli tips. Each line represents data collected from 20 z-plane images of a cell; the red line is the mean of the 19 plots. (J) Normalized GP values for each line in (I). (K) The mean *SR*’ values between the CD45 and IgM images segmented by distance obtained from 19 cells were plotted as a function of the distance from microvilli tips. Error bars represent the SD.

### Depletion of cholesterol diminishes the pre-exclusion of CD45 from the tips of microvilli

We hypothesized that the accumulation of cholesterol at the microvilli tips causes local thickening of the PM at the tips, which in turn, creates a diffusion barrier for CD45. To test this hypothesis, we took two independent approaches: depleting cholesterol from the cell membrane using MβCD (Mahammad and Parmryd, 2008) and engineering the CD45 transmembrane domain.

Freshly isolated (resting) or effector human CD4^+^ T cells were treated with 0, 5, or 10 mM MβCD for 30 min to extract cholesterol from the membrane (Figures 4A-F and S6C-G). Treating with 10 mM of MβCD was reported to disrupt lipid rafts in human Jurkat T cells (Larbi et al., 2004). More than 90% of both resting and effector CD4^+^ T cells were still viable after treatment with 10 mM MβCD (Figure S6A). MβCD treatment had no significant effect on the intensity of CD45 and CD3 (Figure S6B). However, the GP values near the microvilli tips increased upon cholesterol depletion, resulting in a decrease in the slopes of the GP distribution plots as a function of higher MβCD concentrations (Figures 4C, 4E, S6D, and S6F). This indicates that the CD45 exclusion from microvilli tips is mitigated by MβCD. Consistent with this, the correlation values between CD45 and CD3 near the tip area calculated by *SR’* became less negative in the MβCD-treated resting and effector cells (Figures 4D, 4F, S6E, and S6G). Note that the *SR’* values at the microvilli tips were still negative in the MβCD-treated samples, indicating some remaining CD45 exclusion, which likely reflects incomplete depletion of cholesterol. The Pearson’s correlation coefficients, *R’* (Table S1), between CD45 and CD3 were also significantly increased in the treated cells (Figures 4G and 4H). These results are consistent with the notion that cholesterol depletion diminished CD45 pre-exclusion from the microvilli tips.

**Figure 4.**
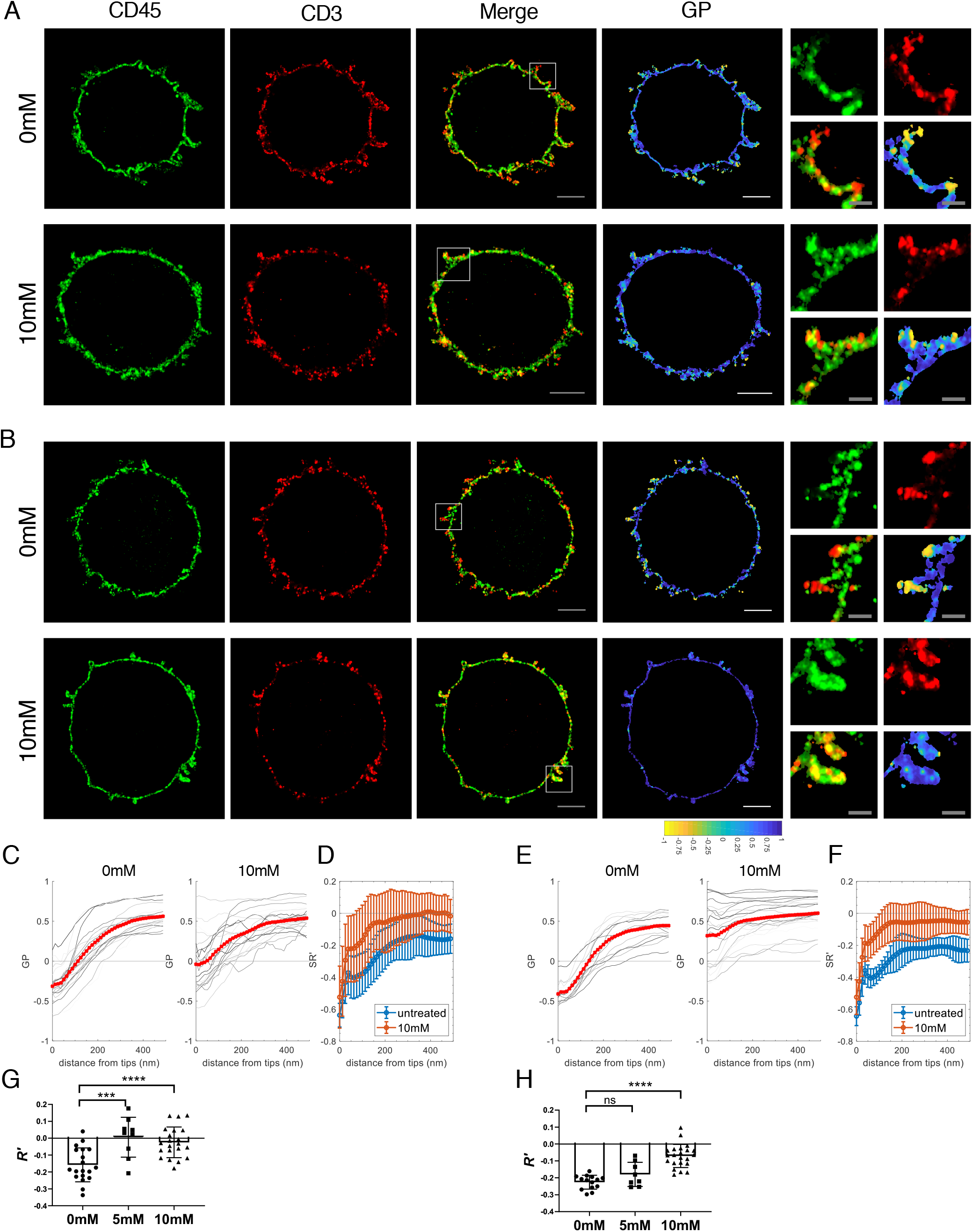
Effect of MβCD on the distribution of surface CD45 molecules in human CD4^+^ T cells. (A, B) Representative 4x-ExM-Airyscan images of CD45 (green), CD3 (red), the merged, and GP of a representative untreated (0 mM) or MβCD-treated (10 mM) resting (A) and effector (B) T cells. Magnified images of the area marked in the merged images are presented in the right most panels. Scale bars: 2 μm in the merge and the GP images; 500 nm in the magnified images. (C) The median GP values of MβCD-treated resting T cells. 19 cells (0 mM) and 21 cells (10 mM) were plotted as a function of the distance from the microvilli tips. Each line represents data collected from 20 z-plane images of a cell. The red lines are the mean of the plots. (D) The mean *SR’* values between the CD45 and CD3 images of the cells in (C) segmented by distance were plotted as a function of the distance from the microvilli tips. Error bars represent the SD. (E, F). The median GP values and the mean *SR’* values between the CD45 and CD3 of MβCD-treated effector T cells. 14 cells (0 mM) and 22 cells (10 mM) were analyzed as in (C) and (D). (G, H) The mean Pearson’s correlation coefficient (*R’*) of CD45 and CD3 images of untreated, 5 mM, or 10 mM MβCD-treated resting (G) or effector (H) T cells. ***, *P* < 0.001; ****, *P* < 0.0001; ns, not significant, calculated by the unpaired Student’s t-test.

### A short membrane integration limit (MIL) of the CD45 transmembrane (TM) domain accounts for its pre-exclusion

We considered the possibility that the relatively short “membrane integration limit” (MIL) of CD45 may account for its exclusion from the thicker microvilli tips. We define MIL as the maximum number of amino acids (a.a.) between two charged residues (“hard stops”) located nearest to each end of the TM domain of a PM-residing protein (Table S2), which defines the maximum sequence length that can potentially be integrated into the PM. These “hard stop” charged residues include lysine (K), arginine (R), glutamic acid (E), aspartic acid (D), and histidine (H). For instance, as shown in Figure 5A, the predicted TM domain lengths of both CD45 and CD3δ are 21 a.a. However, the MIL of CD45 is 22 a.a., flanked by two positively charged lysine (K) residues at each end, while that of CD3δ is 27 a.a. between aspartic acid (D) and histidine (H). We define the “extra” a.a. residues within the MIL outside the TM domain as “spacer”. Since charged a.a. are most unlikely to be integrated into the hydrophobic core in the bilayer (Engelman et al., 1986; Kyte and Doolittle, 1982), this MIL represents the maximal number of a.a. (“spacer” plus TM domain) that can theoretically be integrated into the membrane.

**Figure 5.**
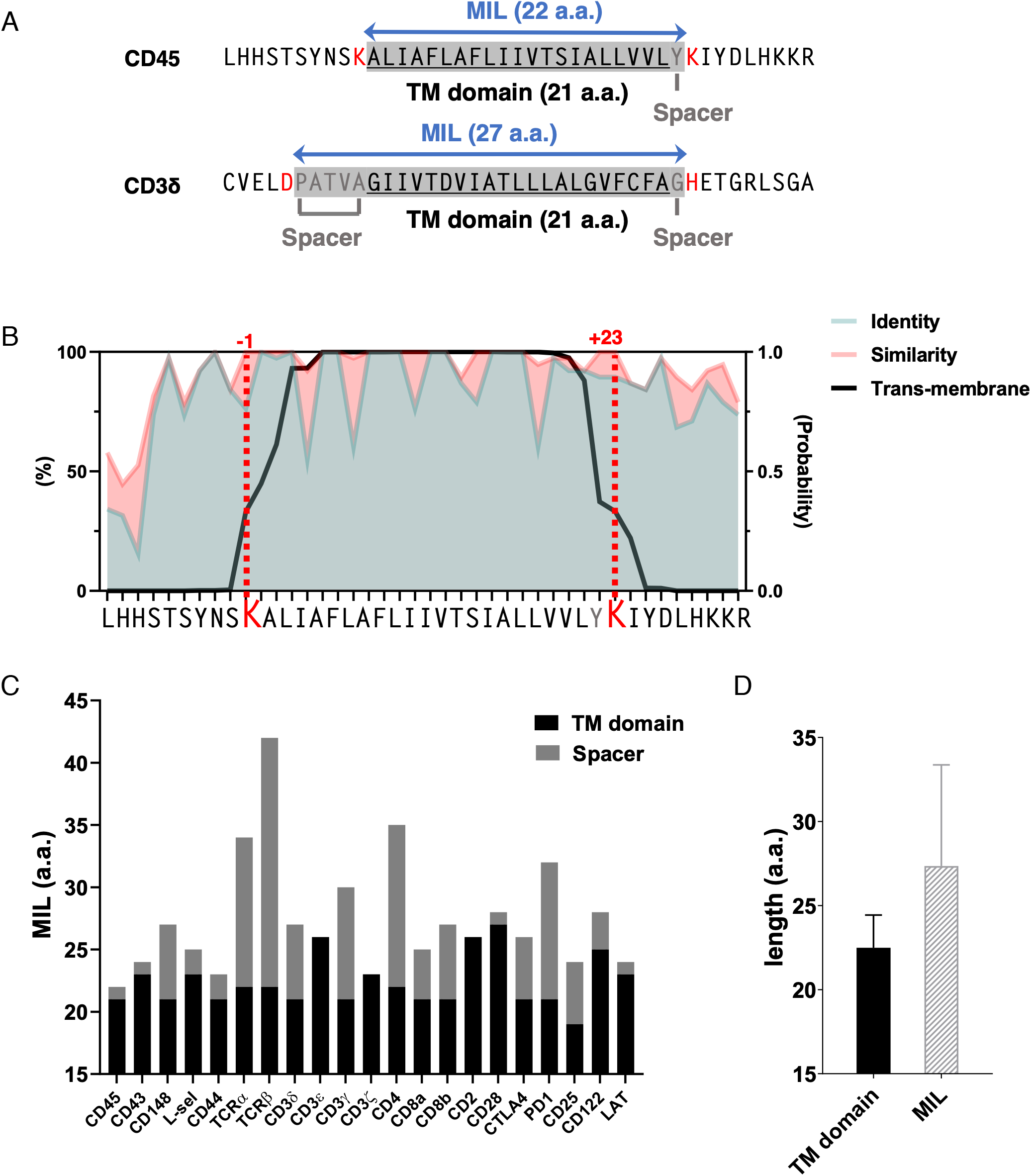
Analysis of TM domain and MIL lengths. (A) Examples of the MIL of human CD45 and CD3δ. Shown are the a.a. sequences of TM domain (underlined) ±10 flanking residues. The charged a.a. residues located at each end of the TM domain are marked in red; spacers (residues between the TM domain and the charged a.a. residues) are marked in gray; The MIL are marked by blue double arrows. (B) The percentages of sequence identity or similarity (allowing substitutions within the same group of a.a.: charged [K, H, R, E, D]; hydrophobic [A, I, L, F, V, P, G]; polar [Q, N, S, T, C]; amphipathic [W, Y, M]), and the probability of TM positioning of residues analyzed using TMHMM v2.0 (transmembrane helices based on a hidden Markov model (Sonnhammer et al., 1998) (http://www.cbs.dtu.dk/services/TMHMM/) for each residue of the CD45 TM domain ± 10 flanking residues among 38 species (Table S3). (C) Lengths of the TM domains (black bars) and spacers (gray bars) of the T cell membrane proteins listed in Table S2. (D) Mean lengths of the TM domains and the MILs of non-CD45 membrane proteins listed in Table S2. Error bars represent the SD.

The charged a.a. residues at position K^577^ and K^600^ of human CD45 are 100% conserved among 38 different species (Figure 5B, Table S3), and the MIL of these CD45 molecules (22 a.a.) is shorter than that of most other major TM proteins expressed on T cells, B cells, or APCs (Figures 5C, 5D, and Table S2). The mean MIL length between charged a.a. (K, R, E, D, H) of the non-CD45 membrane proteins (Table S2) is 27.4 a.a, which is predicted to be a 41.1 Å-long α-helix [1.5 Å per residue (Pauling et al., 1951)], while the 22 a.a. MIL of CD45 is predicted to be only a ~33Å-long α-helix, which matches the thickness of a pure phospholipid bilayer, but is substantially shorter than the thickness of the membrane at lipid raft domains. Therefore, we hypothesized that CD45, which has a shorter MIL, cannot diffuse into the thicker membrane at the microvilli tips, resulting in its exclusion, while other proteins that have a longer MIL can.

To test whether the shorter MIL of CD45 and/or its long extracellular domain were responsible for its pre-exclusion from the microvilli tips, we constructed two CD45 mutants with an added N-terminal triple hemagglutinin A (HA) tag sequence (Figure 6A): CD45ΔEC, from which the extracellular domain of human CD45 was deleted except 6 a.a. adjacent to the TM domain; and CD45ΔECMIL25, in which the lysine (K) at position +23 in the C-terminus of the TM domain was replaced with a hydrophobic a.a., leucine (L) of the CD45ΔEC, thereby increasing its MIL from 22 in the wild-type CD45TM domain to 25 a.a. (Figure 6A). We then determined the distribution of these mutant CD45 molecules and compared it to that of endogenous CD45 in lentivirally transduced Jurkat T cells by staining with anti-HA or anti-CD45 antibodies, respectively. The cells properly expressed their endogenous CD45, as well as the transduced CD45 mutant (Figures 6B, 6C, S7, and S8).

**Figure 6.**
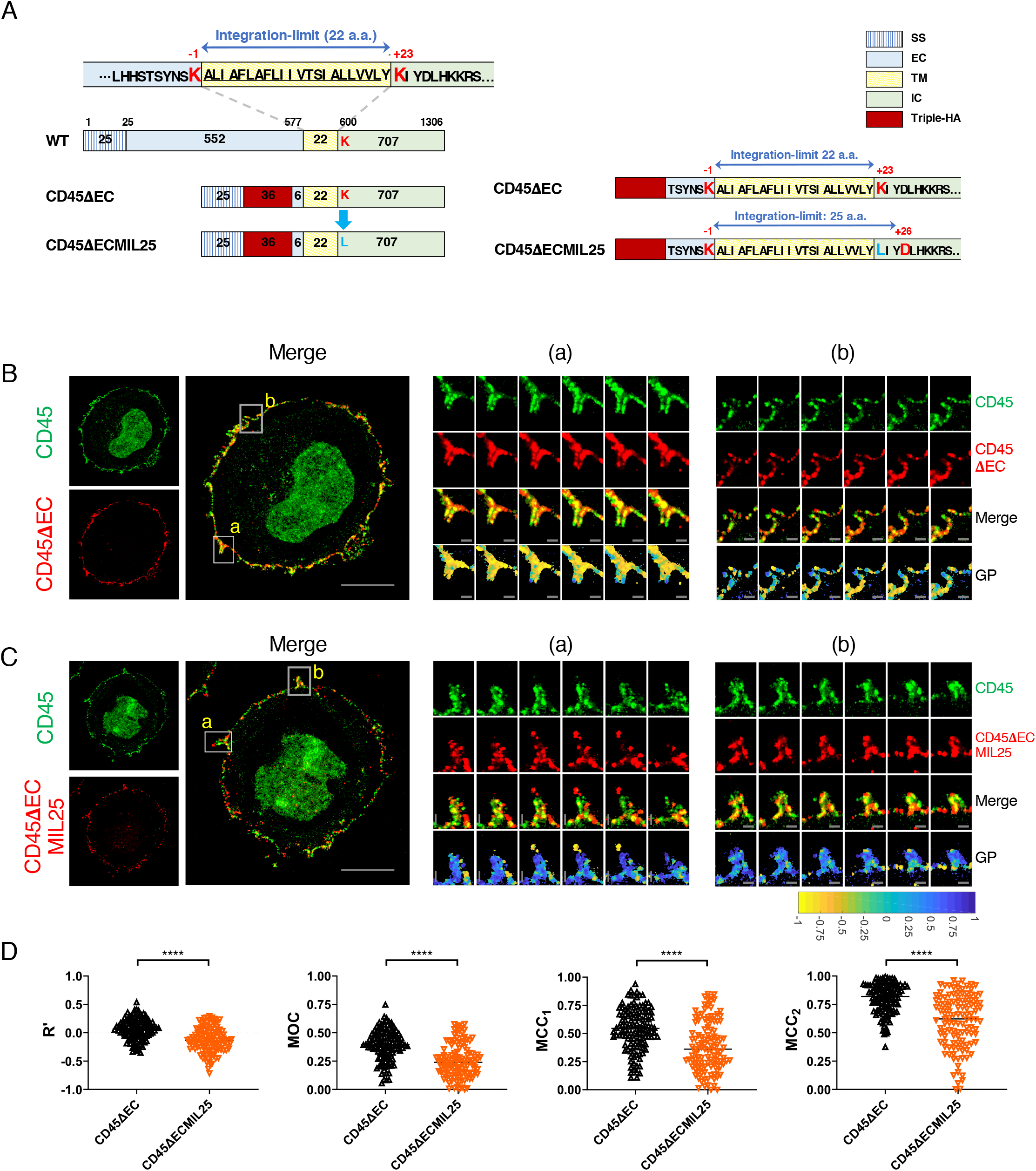
CD45 distribution in Jurkat T cells lentivirally transduced with CD45 mutants. (A) A schematic representation of wild-type (WT) human CD45 gene and CD45 mutants, CD45ΔEC and CD45ΔECMIL25. CD45ΔEC (native TM domain) and CD45ΔECMIL25 (K600L) include the signaling sequence domain (SS), a short extracellular domain (6 a.a.) (EC), TM domain (TM), intracellular domain (IC) and triple HA tag sequences. MIL of CD45ΔEC and CD45ΔECMIL25 are marked on the right with blue double arrows. (B, C) Representative 4x-ExM-Airyscan images of Jurkat T cells lentivirally transduced with CD45ΔEC (B) and CD45ΔECMIL25 (C) CD45 mutants. Images of endogenous (green, upper left) or the transduced (red, lower left) CD45, and the merged images between the two are presented. Scale bars: 5 μm. Six z-stack images (step size 125 nm) of the areas marked (a) and (b) in the merged images were magnified in the right panels. Scale bars: 500 nm. The color bar at the bottom represents GP values between −1 (100% mutant expression) and +1 (100% endogenous CD45 expression). (D) Colocalization analysis between the endogenous and mutant CD45 of 120 microvilli images (Figure S8) collected from each CD45ΔEC (black)- and CD45ΔECMIL25 (orange)-transduced Jurkat cells. Pearson’s correlation coefficient (*R’*), Manders’ overlap coefficient (MOC), Manders’ correlation coefficients (MCC_1_ and MCC_2_) were calculated as described in Table S1 and Method. ****, *P* < 0.0001, calculated by the Mann-Whitney test.

The CD45ΔEC-transduced cells displayed clear exclusion of the mutant CD45 similar to that of the endogenous CD45 (Figures 6B, S7A, and S8A), indicating that CD45 pre-exclusion from the tips of microvilli was not driven by the long extracellular domain. In sharp contrast, the CD45ΔECMIL25-transduced cells no longer displayed exclusion of the mutated CD45 from the microvilli tips (Figures 6C, S7B, and S8B). The negative GP values found at the microvilli tips indicated that the CD45ΔECMIL25 mutant was neither restricted to nor excluded from the microvilli tips. Because the expression levels of both CD45ΔECMIL25 and endogenous CD45 were low and discontinuous on the surface, we had to modify the analysis method from that used in primary cells (*i.e.,* Figures 2H-K). We selected 120 microvilli images each collected from the CD45ΔEC- or CD45ΔECMIL25-transduced cells (Figure S8) and analyzed the Pearson’s correlation coefficient (*R’*), Manders’ overlap coefficient (MOC), and Manders’ correlation coefficients (MCC_1_ and MCC_2_) (Table S1) (Manders et al., 1993) between endogenous and mutant CD45. As shown in Figure 6D, those colocalization coefficients were significantly reduced in CD45ΔECMIL25-transduced cells compared to CD45ΔEC-transduced cells. Thus, we conclude that the increased MIL of the TM domain in the CD45ΔECMIL25 mutant, rather than the extracellular domain, enabled this CD45 mutant to diffuse into the membrane of microvilli tips and, therefore, that the short MIL of the native, endogenous CD45 TM domain is indeed responsible for pre-exclusion from the microvilli tips.

### Early microvilli-mediated interactions between T cells and APCs induce Ca^2+^ influx

To determine the functional significance of CD45 pre-exclusion, we tested whether microvilli-mediated transient contacts with APCs can trigger TCR signaling prior to the formation of a stable IS, at which time CD45 exclusion is driven by size exclusion from the tight T cell-APC contacts (Chang et al., 2016; Razvag et al., 2018). We predicted that the absence of (or lesser) inhibition of TCR signaling by CD45 at the tips of microvilli at this early time due to CD45 pre-exclusion may enable or facilitate local Ca^2+^ triggering from brief T cell-APC contacts before a stable, flattened IS is formed. T cell microvilli are known to scan the APC surface (Cai et al., 2017), but it was not known whether these transient microvilli-mediated contacts can induce Ca^2+^ influx in T cells. For this analysis, we applied fast sub-diffraction limit 3D optical diffraction tomographic (ODT) live cell microscopy (Kim et al., 2013). We used an antigen-specific system, OTII x *Rag2*^−/−^ (KO) TCR-transgenic CD4^+^ T cells specific for ovalbumin (OVA), which were loaded with the Ca^2+^ indicator Fluo-4 AM and stimulated with OVA peptide-pulsed B cell APCs. The maximum theoretical lateral resolutions of the ODT system can reach ~110 nm (Lauer, 2002; Park et al., 2020). The practical lateral resolution of the system was reported to be 172 nm (Shin et al., 2018). As shown in Video S1 and S2, T cells established contacts with antigen-presenting B cells (OVA-pulsed) via their microvilli, and these brief contacts induced Ca^2+^ flux in T cells. The Ca^2+^signals often fluctuated during the microvilli-mediated contacts. Thus, the Ca^2+^ flux oscillations previously observed during initial T cell-APC contacts (Le Borgne et al., 2016) were likely due to the brief early contacts made by microvilli, which could not be resolved in the previous study.

## Discussion

Here we report a novel feature of T cell microvilli, namely, CD45 pre-exclusion from their tips prior to engagement by APCs. This pre-exclusion was observed in human and mouse lymphocytes, CD4, CD8 and regulatory T cells, and B cells. This spatial pre-exclusion of CD45 resulted in CD45-free TCRs on the microvilli tips. The CD45 pre-exclusion is caused by the short MIL (22 aa) of the CD45 TM domain that prevents CD45 from diffusing into the thicker PM at the tips of microvilli due to the higher cholesterol content at its negative curvature.

It has been reported that CD45 exclusion from the IS was initiated from isolated tight contacts originating from microvilli (Chang et al., 2016; Razvag et al., 2018). Here, we show for the first time that CD45 exclusion is actually present before forming a tight contact with APCs. A theoretical model of TCR triggering suggested that two factors are key to initiating TCR signaling: (i) the dwell time of TCRs in the CD45-free area; and (ii) spatial constraints on contact area size imposed by cell topography (Fernandes et al., 2019). Our findings that TCRs are present in the CD45-free zone at small constrained areas of the microvilli tips is consistent with the model accounting for how TCR sensitivity is maintained while avoiding non-specific activation. CD45 pre-exclusion has distinct consequences: 1) The absence of “long” molecules at or near the tip of microvilli facilitates scanning and engagement of APCs presenting pMHC complexes; and 2) The absence of CD45 phosphatase activity enables effective initiation of TCR signaling.

CD45 pre-exclusion discovered herein suggests a new facet of early TCR signaling, *i.e.*, the ability of CD45-free TCRs at the tips of microvilli to effectively scan the surface of APCs and transduce signals. This clearly happens prior to the formation of a mature IS and may nucleate CD45 exclusion from the IS, which progresses as the T cell-APC contact area is enlarging. During the IS formation process, collapsing of the microvilli (Ueda et al., 2011) may allow the additional TCR molecules to be recruited from the columns of microvilli to the IS and the CD45 in the column to be exclude by size exclusion due to their long extracellular domain from the 2D-flattened interface (Schmid et al., 2016; Shaw and Dustin, 1997).

Our study is the first to demonstrate that Ca^2+^ influx is induced at the microvilli-mediated contact areas prior to the formation of a mature, flattened IS. Recent studies showed that CD45 segregates from isolated tight microvilli contacts within seconds of TCR engagement, and that TCR phosphorylation negatively correlates with this TCR-CD45 separation (Razvag et al., 2018; Razvag et al., 2019). Moreover, a recent study showed that the TCR coreceptors, CD4 and CD2, and early signaling molecules, Lck and LAT, are highly enriched on microvilli (Ghosh et al., 2020). Together, these findings implicate microvilli as specialized cellular domains where initial antigen recognition and signal transduction occur. Thus, TCRs on the tips of microvilli are ready to trigger upon engagement of pMHC, whereas TCRs on the columns of microvilli need to be segregated from CD45 during IS formation.

We hypothesized that the molecular mechanism on CD45 pre-exclusion may be related to the curvature-driven lipid partitioning of the PM at the tips of microvilli and the limited length of the CD45 TM domain. It has been suggested that the highly curved membrane at the tips of microvilli can lead to local sorting of specific lipids (Cebecauer et al., 2018). Although the specific lipid composition at the tips of the protrusive membrane regions is unknown, studies in model membranes have shown that cholesterol accumulation at highly negative curvatures promotes membrane stability (Wang et al., 2007). A recent analysis of lipid composition on a cell membrane using nanoscale secondary ion mass spectrometry (nanoSIMS) imaging showed that cholesterol-enriched sphingolipid patches highly accumulated at substrate-bound cellular projections coinciding with the tips of filopodial protrusions (Yeager et al., 2018). This finding supports our hypothesis that cholesterol-rich domains exist at the tips of microvilli.

We first tested our hypothesis by using MβCD to deplete membrane cholesterol in resting and effector T cells, and found that this treatment reduced the exclusion level of CD45 from the microvilli tips. We also observed that the microvilli in MβCD-treated cells were fewer and thinner compared to the ones in untreated cells, especially in resting CD4^+^ T cells. A similar reduction in the number of microvilli was reported in MβCD-treated MDCK cells and B cells (Greicius et al., 2004; Poole et al., 2004), indicating that cholesterol promotes the stability of microvilli. Secondly, using Jurkat T cells lentivirally transduced with two different CD45 mutants, we ruled out a role for the extracellular CD45 domain in pre-exclusion. This is consistent with a recent report that the CD45 TM domain lacking most of the extracellular domain and the intracellular domain was segregated out of the ordered (raft) domain in the membrane, where BCRs were enriched in the raft domain (Stone et al., 2017). Importantly, we demonstrated that increasing the MIL of the CD45 TM domain by replacing the lysine (K) residue in the +23 position with leucine (L) allowed the CD45ΔECMIL25 mutant to diffuse into the microvilli tips.

In conclusion, the pre-existing CD45 exclusion from microvilli tips discovered here strongly supports a novel three-phase model of formation of the IS. First, the absence of CD45 phosphatase activity at the tips of microvilli represents small phosphatase-free zones at the tips of microvilli. This initiates TCR triggering on the tips of microvilli that engage a single or very few antigen-pMHC complexes while scanning the surface of an APC. Second, microvilli are collapsed, resulting in more TCRs from the columns of microvilli engaging pMHC, and the CD45 molecules situated within the flattened interface are driven out from the IS because of size exclusion of their extracellular domain. Third, a mature IS is formed and fully establishes a T cell response (Kumari et al., 2019).

## Acknowledgments

This is manuscript number 3383 from the La Jolla Institute for Immunology (LJI). We thank Drs. Sara McArdle (La Jolla Institute for Immunology), Enrico Gratton (University of California, Irvine), and Gilad Haran (Weizmann Institute of Science) for helpful discussions on imaging analysis, Dr. Padmini Rangamani and Mr. Arijit Mahapatra (UCSD) for discussions on membrane modeling, Ms. Kristine Suchey (Division of Laboratory Animal Care, LJI) for collecting mice tissues, and members of the microscopy and flow cytometry cores (LJI) for technical support. We appreciate CTK Instruments and Mr. Adam Lucio for offering us to use the Tomocube system and for the technical support. This research was supported by NIH grants S10OD021831 and P01 HL078784 project 3 to KL

## Author Contributions

Conceptualization, Y.J.; Methodology, Y.J., and L.W.; Software, Y.J. Formal Analysis: Y.J. Investigation, Y.J., Resources, K.L.; Data Curation, Y.J. Writing – Original Draft, Y.J.; Writing – Review & Editing, Y.J., A.A, and K.L; Visualization, Y.J.; Supervision, K.L and A.A.; Funding Acquisition, K.L.

## STAR Methods

### RESOURCE AVAILABILITY

Further information and requests for resources and reagents should be directed to and will be fulfilled by the Lead Contact, Yunmin Jung.

#### Materials Availability

Plasmids generated in this study are available upon request.

#### Data and Code Availability

The supporting data and the Matlab code used in this study are available from the lead contact upon request.

### EXPERIMENTAL MODEL AND SUBJECT DETAILS

#### Mice

C57BL/6J mice were purchased from Jackson Laboratory. *Foxp3*^YFP-Cre^ reporter mice were kindly gifted by Dr. Lynn Hedrick (La Jolla Institute for Immunology). OT-II x *Rag2*^−/−^ mice were generated by crossing male OT-II x *Rag2*^−/−^ mice kindly gifted by Dr. Stephen Schoenberger (La Jolla Institute for Immunology) with female *Rag2*^−/−^ mice obtained from Taconic. Mice were maintained and used by following the guidelines of the La Jolla Institute for Immunology Animal Care and Use Committee, and approval for use of mice was obtained from the La Jolla Institute for Immunology according to criteria outlined in the Guide for the Care and Use of Laboratory Animals from the National Institutes of Health.

#### Cells

Human CD4^+^, CD8^+^, and B cells were purified from peripheral blood mononuclear cells (PBMCs) prepared from the whole blood of healthy donors through the La Jolla Institute for Immunology Clinical Core using EasySep negative selection kits (StemCell Technologies). Freshly isolated resting cells were used after incubation in complete phenol red-free RPMI (Gibco) growth medium [2 mM L-glutamine (Gibco), 1 mM sodium pyruvate (Gibco), 1 mM nonessential amino acids (Gibco), 100 U penicillin-streptomycin (Gibco), and 10% (v/v) FBS (Gemini Bio)] for ~2-4 hours in a CO_2_ (5%) incubator at 37°C. For effector T cells, freshly isolated CD4^+^ T cells were stimulated on an anti-CD3 plus anti-CD28 antibody-coated plate for 2 days and then transferred and grown in a complete medium supplemented with IL-2 (100-130 U/mL) and 2-mercaptoethanol (50 μM) for 7 days. Mouse spleens and lymph nodes were processed through a 70 μm filter (BD Biosciences) and the single cell suspensions were washed and incubated with RBC lysis buffer (eBioscience) for 5 min at room temperature (RT). After washing, CD4^+^ T cells or B cells were purified using EasySep negative selection kits. The purified cells were used after a short incubation at 37°C as described above for human cells. For ODT live imaging, ovalbumin-specific CD4^+^ T cells were isolated from spleens and lymph nodes of TCR-transgenic OT-II mice that were crossed to *Rag2*^−/−^ mice, and mouse B cells used as APCs were isolated from spleen and lymph nodes of gender and age matched C57BL/6J mice using the EasySep negative selection kits. Cells were incubated in complete RPMI growth medium for more than 1 hour in a CO_2_ incubator at 37°C after the purification. Jurkat T cells were grown in phenol red-free RPMI growth media containing 2 mM L-glutamine, 100 U penicillin-streptomycin and 10% FBS. 293T cells were grown in DMEM medium (Invitrogen) containing 10% FBS and 100 U penicillin-streptomycin.

### METHOD DETAILS

#### Antibody labeling

Mouse anti-human CD62L (L-selectin; DREG-56) were labeled with AF568 using an AF568 antibody labeling kit (Thermo Fisher). Purified mouse monoclonal antibodies specific for human CD3 (UCHT1), HA.11 epitope tag (16B12), human IgM (MHM-88), and human IgD (IA6-2) or rat anti-mouse CD62L (MEL-14) were labeled with CF633 using Mix-n-Stain™ CF™ 633 antibody labeling kit (Sigma).

#### Sample preparation for imaging analysis

Human cells suspended in cold PBS (Gibco) containing 5 mM EDTA (Sigma) were washed by centrifugation at 4°C, resuspended in blocking buffer [PBS containing 2% BSA, 5 mM EDTA, 10 mM EGTA, and 0.05% N_3_Na], and incubated on ice for 10 min. Fluorescently labeled antibodies (10 μg/ml) were added to the cell suspension and incubated for additional 20 min on ice. Cells were washed 2x with 5 mM EDTA/PBS and fixed with fixation buffer [4% (w/v) paraformaldehyde (PFA) and 0.5% (w/v) glutaraldehyde (GA; both from Electron Microscopy Sciences), 2% (w/v) sucrose, 10 mM EGTA, 5 mM EDTA, and 0.05% (w/v) N_3_Na (Sigma) in PBS] on ice for 1-2 hours. Cells were washed 2x with PBS, resuspended in cold PBS and kept on ice. For MβCD treatment, cells were washed 2x with serum-free RPMI at RT and incubated without (0 mM) or with 5 mM or 10 mM of MβCD (Sigma) in RPMI containing 25 mM HEPES (Sigma) in a CO_2_ incubator for 30 min at 37°C. Cells were washed 2x with cold 5 mM EDTA/PBS at 4°C, and then incubated in blocking buffer containing 2% (w/v) fatty acid-free BSA (Sigma), 5 mM EDTA, 10 mM EGTA, and 0.05% N_3_Na in PBS. For mouse samples, mouse T cells and B cells were prepared as described above except cells were resuspended, washed, and fixed in cytobuffer [2 mM MgCl_2_, 5 mM EGTA, and 2% sucrose in 80 mM HEPES]. For membrane labeling, the cells were stained with FM 4-64FX (Molecular Probes) after overnight incubation for the 4x-ExM sample preparation (see the 4x-ExM-Airyscan microscopy section). After washing with PBS at RT, cells were stained with FM 4-64FX (5 μg/ml; Invitrogen) in PBS for 1 hour at RT. Cells were washed 3x with PBS, and then fixed and washed as described above. For ODT live imaging, isolated OT-II T cells were stained with Fluo-4 AM (2.5 μM; Molecular Probes) and incubated in a CO_2_ incubator at 37°C for 30 min. C57BL/6J mouse B cells, used as APCs, were washed and resuspended in serum-free, phenol-red-free RPMI and were incubated with 10 μg/ml of OVA_323-339_ peptide (Invivogen) at 37°C for 1 hour. After 30 min, Cell Tracker Orange CMRA Dye (0.5 μM; Molecular Probes) was added to the cells. Both OT-II T cells and B cells were washed twice and resuspended in imaging buffer [48% phenol red-free RPMI, 50% HBSS (Gibco), 2% FBS].

#### Plasmids and lentiviral transduction

DNA sequences encoding the CD45 mutants, CD45ΔEC and CD45ΔECMIL25, were synthesized (Integrated DNA Technologies) and cloned into the lentiviral vector pFUW (Lois et al., 2002) (a gift from Alok Joglekar and David Baltimore). The resulting vectors pFUW-CD45ΔEC or pFUW-CD45ΔECMIL25 were confirmed by DNA sequencing. The two vectors along with two packaging plasmids pSPAX2 and pMD2.G were cotransfected into 293T cells using the TransIT-LT1 Transfection Reagent (Mirus Bio LLC) according to the manufacturer’s instructions. The supernatant was collected after 48 hours, and applied to human T cells for viral delivery. Two days later, human T cells were harvested for microscopic or for flow cytometric analysis.

#### Flow cytometry and sorting

Some of the MβCD-treated cells were collected from the blocking reaction to determine viability by diluting them with PBS and staining with 7-AAD (20 μg/ml, BD Biosciences) for 20 min on ice, followed by washing with cold 1% FBS/PBS. The cells were resuspended in 200 μl of 1% FBS/PBS and kept on ice in the dark. Cells were immediately loaded for flow cytometry acquisition on a BD FACSCelesta (BD Biosciences). Single cells gated using forward *versus* side scatter (FSC *vs*. SSC) were analyzed for 7-AAD fluorescence using FACSDiva software (BD Biosciences). Data analysis was done with FlowJo software. For sorting the Treg cells, CD4^+^ cells isolated from *Foxp3*^YFP-Cre^ mice were labeled with Alexa Flour 488-conjugated anti-mouse CD45 and CF633-conjugated anti-mouse L-selectin on ice for 20 min as described in sample preparation for mouse cells. Fixed cells were resuspended in 0.5 ml PBS and were kept at 4°C until sorting. After selecting single cells determined by FSC *vs*. SSC gating, CD45^+^Lsel^+^YFP^+^ cells were sorted and collected in PBS on a BD Aria III or Fusion cell sorter (BD Biosciences). Jurkat T cells transduced with CD45 mutants CD45ΔEC, CD45ΔECMIL25 were washed 2x in serum-free RPMI at RT. Alexa Flour 488-conjugated anti-human CD45 and CF633-conjugated anti-HA were labeled in 2% FBS/PBS for 20 min at RT. After washing twice, cells were fixed with 4% (w/v) PFA, 0.5% GA, 2% sucrose, 10 mM EGTA, 5 mM EDTA and 0.05% N_3_Na in PBS on ice for 1 hour at RT. Cells were washed twice in PBS and resuspended in 0.5 ml PBS. After selecting single cells determined by FSC *vs*. SSC gating, CD45^+^ cells expressing high level of HA were sorted and collected in PBS on a BD Aria III or Fusion cell sorter (BD Biosciences). The sorted cells were processed for 4x-ExM-Airyscan imaging as described below.

#### 4x-ExM-Airyscan imaging

The 4x-ExM samples were prepared as described in (Chen et al., 2015; Tillberg et al., 2016) with some modifications. Succinimidyl ester of 6-((acryloyl)amino)hexanoic acid (0.1 mg/ml; acryloyl-X, SE, Life Technologies) was added to the fixed cells in PBS and then the 35 μl of cells were placed on freshly prepared poly-L-lysine (PLL; 0.1%; Sigma)-coated 5 mm round glass coverslips (Electron Microscopy Sciences). Samples were kept overnight at 4°C, and rinsed 4x with PBS at RT. Buffer was removed by gentle suction and then the cell-coated coverslip was quickly placed on top of a second rectangular glass coverslip (15 ×15 mm; Electron Microscopy Sciences) that had a drop of freshly mixed gelation solution. The gelation solution was freshly mixed with monomer solution [PBS supplemented with 2 M NaCl, 8.625% (w/w) sodium acrylate, 2.5% (w/w) acrylamide, 0.15% (w/w) N,N′-methylenebisacrylamide, ammonium persulfate initiator (APS; 0.2%; Sigma) and tetramethylethylenediamine (TEMED; 0.2%; Sigma)], kept on ice before use. Gelation samples were sealed and kept in the dark for 45 min at RT, followed by digestion of the gel in digestion solution [50 mM Tris buffer (pH 8.0) containing 1 mM EDTA, 0.5% Triton X-100, 0.8 M guanidine HCl (Sigma) and 1% (v/v) proteinase K (New England Biolabs)] for 2 hours at 37°C degrees with gentle shaking. Gels formed between the two coverslips were detached during the digestion, fully expanded with an excess of H_2_O for 30 min at RT, and immobilized on PLL-coated (0.1%; Sigma) coverslip for microscopy. Cells were imaged on a laser scanning confocal microscope ZEISS LSM 880 equipped with an Airyscan detector (Carl Zeiss). Airyscan images were taken with a C-Apochromat 40x/1.2 W AutoCorr M27 objective with a 100-130 μm sized pinhole with master gain 850 or 950 while keeping humidity in the sample chamber by a humidifier. The 488-, 561-, or 633-nm laser lines (Carl Zeiss) were used to excite the AF488-, AF568- and CF633-labeled samples, respectively, and a 561-nm laser line was used to excite FM 4-64FX. Doubly labeled samples (AF488- and CF633- or AF568-conjugated antibodies) were acquired with interleaved laser excitation (ILEX) mode. The excitation and emission for each channel were separated with a combination of a filter set of a beam splitter MBS 488/561/633 (Carl Zeiss) and multi bandpass filter BP 495-550 + LP 570 (Carl Zeiss). Triple-channel images were acquired sequentially frame by frame with the main beam splitter MBS 488/561/633 combined with SBS SP 615 (Carl Zeiss) and BP 420-480 + BP 495-550 for AF488-labeled samples, SBS SP 615 and BP 495-550 + LP 570 (Carl Zeiss) for AF568-labeled samples; and SBS LP 660 (Carl Zeiss) and BP 495-550 + LP 570 (Carl Zeiss) for CF633-labeled samples and FM 4-64FX-labeled samples. For each channel, ~80-150 series of z-plane Airyscan confocal images were taken with 0.25 μm steps. The x and y step sizes (a pixel size) were 50 nm. ZEN Black 2.3 SP1 FP2 software (Version 14.0.16.201) was used for the post-3D Airyscan processing with automatically determined default Airyscan Filtering (AF) strength.

#### STORM imaging and analysis

For STORM imaging, μ-Slide 8-well glass bottom chambered coverslips (Ibidi) were cleaned with 1 M NaOH (Sigma) for 1 hour, and coated with PLL (0.1%) for 1 hour at RT before use. Fixed cells labeled with AF647- or AF568-conjugated antibodies and suspended in 300 μl PBS were placed in each well incubated for 1 hour at RT, and then the buffer was exchanged with blinking buffer [PBS containing 0.5 mg/mL glucose oxidase, 40 μg/mL catalase, 10% (w/v) glucose, 50 mM cysteamine, and 93 mM Tris-HCl (all from Sigma)]. All buffers were filtered with 0.2 μm-pore filters (Corning) and freshly mixed before use. For fiducials, Fluorescent Nanodiamond (FND) (100 nm, Adamas Nano) diluted in water (1:1,000) was sonicated for 15 min and then added to the PLL-coated wells for 30 min. Unbound FNDs were washed 5x with distilled water and then blinking buffer was added. STORM imaging was acquired with a UAPON100XOTIRF1.49NA oil objective (Olympus) at epi-illumination mode on Nanoimager (ONI) microscope implemented with a cylindrical lens for localization in z-dimension. The focus was maintained by the autofocus module of the system. Excitation lasers 532 nm and 640 nm and the two channels were separated with dichroic (640 LP) and emission (584/80 and 685/40) filters. 532 nm and 640 nm laser lines were operating at a power of 27.5 kW/cm^2^ and 13.8 kW/cm^2^, respectively. For each channel, 30,000 frames (3,000 frames × 10 movies) were recorded with a speed of 10 ms per frame. The pixel size on the detector was 117 nm. The two-channel registration with FNS fiducial images in the two-channel and single molecule localization processing was performed using Nanoimager software (version 1.1.6165- 012f4ed3; ONI). Data analysis and reconstruction of images were performed with custom-written Matlab (MathWorks) scripts. Single molecule localization data were thresholded as: 0~14 nm for ‘precision in x, y (nm)’; −400~400 nm for ‘z range’; 20,000 for ‘number of photons’; 100 for ‘background’; 0-2.5 for ‘Point spread function σX and σY (pixel)’; 0.6~1.5 for ‘σX/σY’ (pixel)’. Channel correction for individual cell images was done with low-resolution images of the single molecules collected in all frames rendered with the original pixel size by calculating the shifted peak of the 2D cross-correlation image obtained using a 2D Gaussian fit.

#### Optical diffraction tomographic (ODT) microscopy

Freshly prepared OT-II T cells loaded with Ca^2+^ indicator Flou-4-AM and antigen-pulsed B cell APCs labeled with Cell Tracker Orange CMRA Dye were mixed 1:1, and were then immediately loaded onto a TomoDish (Tomocube). Samples on TomoDish were placed on a TomoChamber (Tomocube) to maintain CO_2_ (5%) and temperature (37°C) levels. Cells were subjected to time-lapse imaging with an objective and a condenser lens, UPLASAPO 60XW 1.2NA lens (Olympus), on a holotomographic microscope, HT-2 (Tomocube). Holographic images were generated from interfered images at the camera plane between the two split 532 nm laser beams of a reference beam and a sample illuminated beam obtained at various incident angles modulated by a high-speed illumination scanner using a digital micromirror device (DMD). Following the holographic imaging acquisition in 400 ms, single z-plane fluorescent images of Fluo-4 AM (green channel) for T cells and the Cell Tracker Orange CMRA for B cells (red channel) illuminated with an LED light source (470 nm and 570 nm, respectively) were sequentially acquired with 100 ms exposure time. Time-lapse movies were taken for 5 min with 11-sec intervals for each field of view (37 μm × 37 μm). The refractive index (RI) distribution was reconstructed and visualized for the 3D ODT images using the Tomostudio software (Tomocube). The trace of Fluo-4 AM fluorescence intensities for each frame of the time-lapse movie was analyzed using a custom-written Matlab script. Briefly, green channel image was binarized using the function ‘imbinarize’ (Matlab, 2018a) applying ‘global’ image threshold (using Otsu’s method). Holes in the images were filled with the ‘imfill’ function (Matlab, 2018a). The region of interest (ROI) of a T cell area is determined as of the biggest connected component (with the connectivity of 8) found in the binary image using the ‘bwconncomp’ function (Matlab, 2018a). The mean intensities of the Fluo-4 AM within the ROI were calculated for each frame.

#### Image segmentation

Areas of outside-cell background (OutBG), inside-cell background (InBG), cell body (CB), microvilli (MV) of the 4x-ExM-Airyscan images were segmented using a custom-written Matlab (R2018a, MathWorks) script. The script is available upon request. Briefly, each image channel was converted to binary images (BW1 for channel 1 and BW2 for channel 2) applying the first threshold value obtained using the function ‘multithresh’ in Matlab. The OutBG area was determined as follows: The summed image of the two-channel images was converted to a binary image with a threshold obtained by using the ‘imbinarize’ function in Matlab with an ‘adaptive’ option with the ‘sensitivity’ parameter set to 1. Holes in the binary image were filled with the ‘imfill’ function in Matlab. The OutBG was the area except for the biggest connected component (with the connectivity of 8) found in the binary image using the ‘bwconncomp’ function in Matlab. The InBG area was segmented by ‘activecontour’ function (applying the ‘edge’ method and 25 iterations) in Matlab. Seed mask for the ‘activecontour’ function was generated from dilated images of OutBG. The InBG area was finally obtained after dilated with the function ‘imdilate’ with a disk structuring element with a radius of 5 pixels and holes were filled by ‘imfill’ function. For CB area and MV areas were segmented as follows. Each channel images were converted to binary images using the ‘graythresh’ function with ‘global’ image threshold (using Otsu’s method) and two binary images were added and the non-zero elements of this image were converted to a binary image, and then remove the OutBG or the InBG area in this binary image (SegBW). The GP values, *SR’* coefficients, and intensities for each channel were calculated with data corresponding to SegBW. The SegBW area was then segmented for CB and MV areas for the GP distribution analysis. The CB area was defined by the SegBW area overlapped with the dilated area from the InBG mask image using the ‘imdilate’ function with a disk structuring element with a radius of 24 pixels and the rest of the area in the SegBW was determined as MV area. The locations of the tip of microvilli (microvilli-tip) were determined as follows: The components in the SegBW were connected using imclose function with a disk structuring element with a radius of 5 pixels and then, the small-sized (<2000 pixels) unconnected components were removed for eliminating disconnected microvilli from a cell. The binary image of the MV area in the SegBW was processed with ‘imclose’ (‘disk’, element radius set to 10) and ‘imerode’ (‘cube’, element pixels set to 5) functions, and then were applied to the ‘bwconvhull’ function in Matlab. The output a convex hull image from the ‘bwconvhull’ function was eroded with the ‘imerode’ (‘cube’, element pixels set to 5) function and the zero-elements of this convex hull image overlapped with the MV area were selected for generating a binary image for tips’ areas (TipBW). The TipBW were dilated with the function ‘imdilate (‘cube’, element pixels set to 3) and individual tip-areas (L-BW) were determined with ‘bwlabel’ function with the parameter for ‘connected object size set to 8. The dilated TipBW image was eroded back with the function ‘imerode’ (‘cube’, element pixels set to 3)’ and the mean x,y locations of individual L-BW pixels overlapped with the TipBW were calculated for the location for Microvilli-tips. The length of microvilli was calculated from the closest distance of the location of the Microvilli-tip to an inner boundary of cell determined from the outer-edge pixels of InBG binary image using the ‘edge’ function with ‘Sobel’ method with threshold parameter set to 0 and skipping the edge-thinning stage. Any Microvilli tip location for microvilli whose length was shorter than 50nm or larger than 1.5 μm in human and mouse T cells were excluded for calculation. If any two locations of the Microvilli-tip were closer than 650nm, the Microvilli-tip location for the longer microvilli was excluded for the calculation. The distance from tips of microvilli was calculated using the function ‘pdist2’ in Matlab.

#### Image analysis

Ratiometric generalized polarization (GP), Pearson’s correlation coefficient (*R’*), segment-correlation coefficient (*SR’*), Manders’ overlap coefficient (MOC), and Manders’ correlation coefficients 1 and 2 (MCC_1_, and MCC_2_) values were calculated as shown in Table S1. In Figures 2, 4, S4, and S6, ch1 and ch2 correspond to CD45 and CD3 respectively, in Figures 3, and S5, ch1 corresponds to CD45 and ch2 corresponds to IgM or IgD respectively, and in Figures 6, S7, and S8, ch1 corresponds to the endogenous CD45 and ch2 corresponds to CD45ΔEC or CD45ΔECMIL25 mutants. For colocalization analysis, the first threshold values obtained using the ‘multithresh’ function in Matlab were used for thresholding the positive intensities for each channel. The display range of the intensities of individual color channels in Figure 1 were independently optimized using Zen Black program (Carl Zeiss). The display ranges of intensities in Figure S1C-D (negative) for the L-selectin channel were adjusted to that of Figure S1A for comparison. All the other 4x-ExM-Airyscan images in this study were displayed between the 1% ~ 99.5% intensity of individual channel image using a built-in function in Matlab (MathWorks).

### QUANTIFICATION AND STATISTICAL ANALYSIS

Statistical details of experiments can be found in the figure legends. Mean, median, standard deviation (SD), and the 25^th^ and 75^th^ percentile calculation were performed using the built-in functions in Matlab (MathWorks). Unpaired Student’s t-test and Mann-Whitney test were performed using GraphPad Prism software.

### KEY RESOURCES TABLE

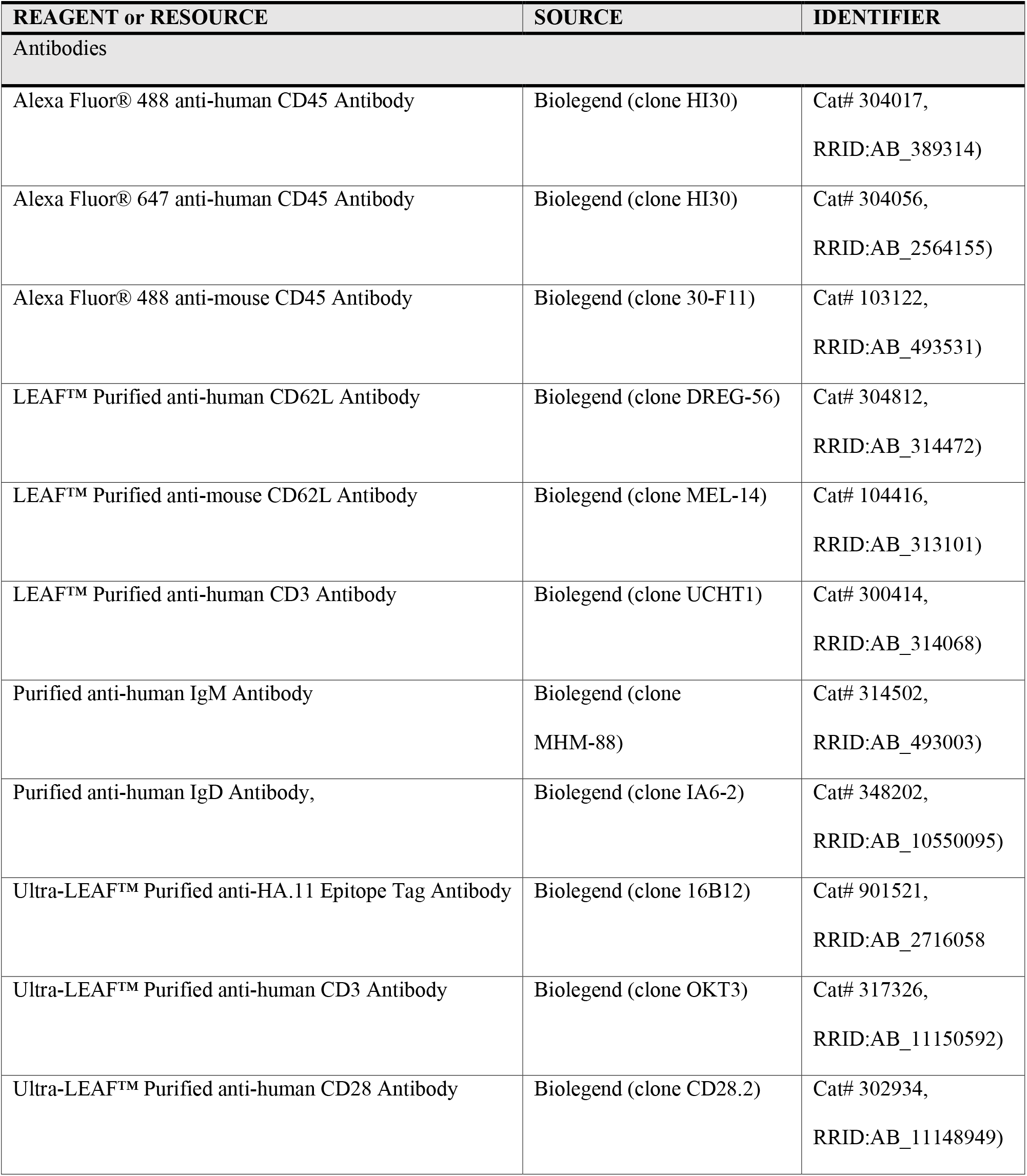
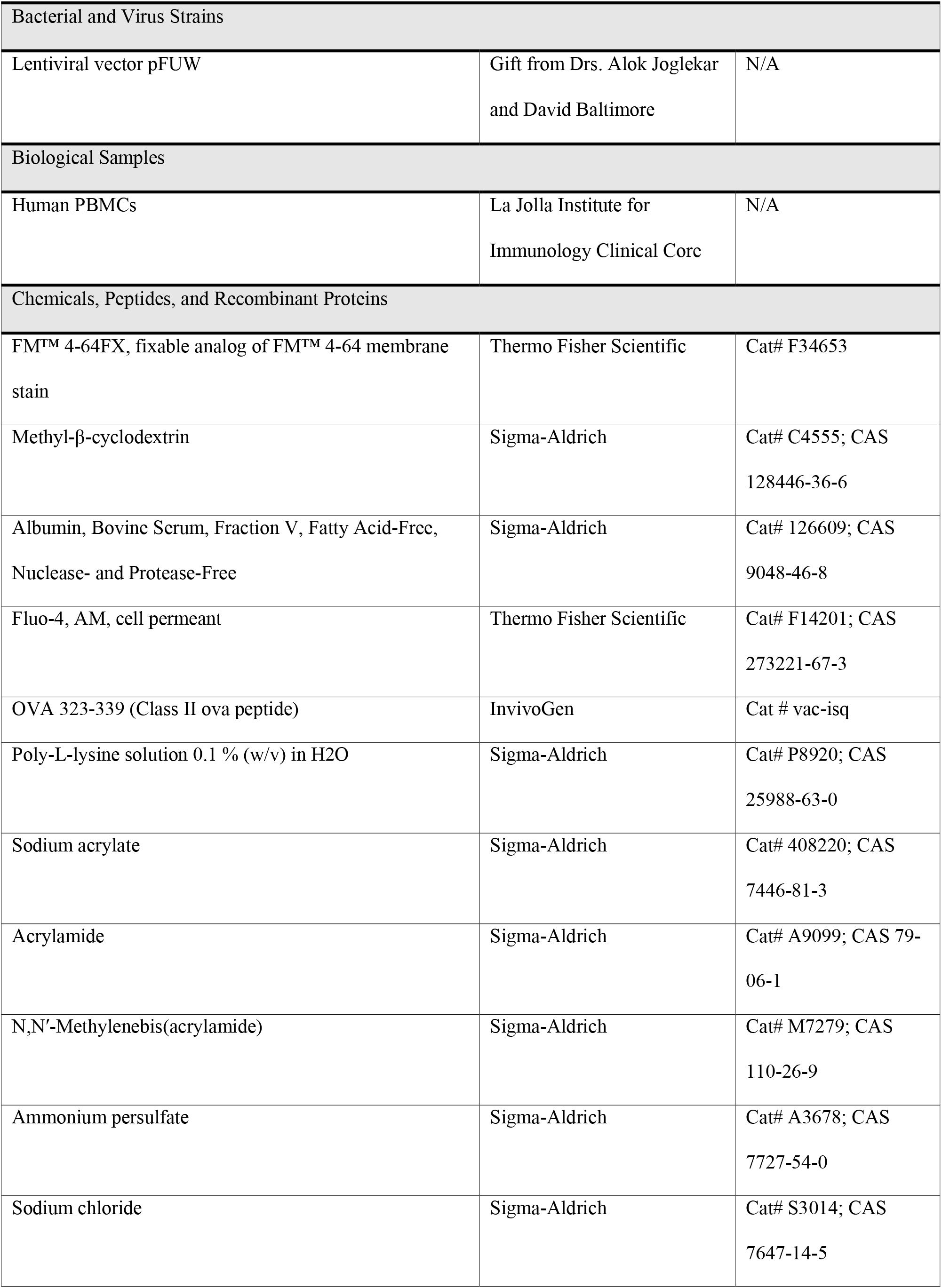
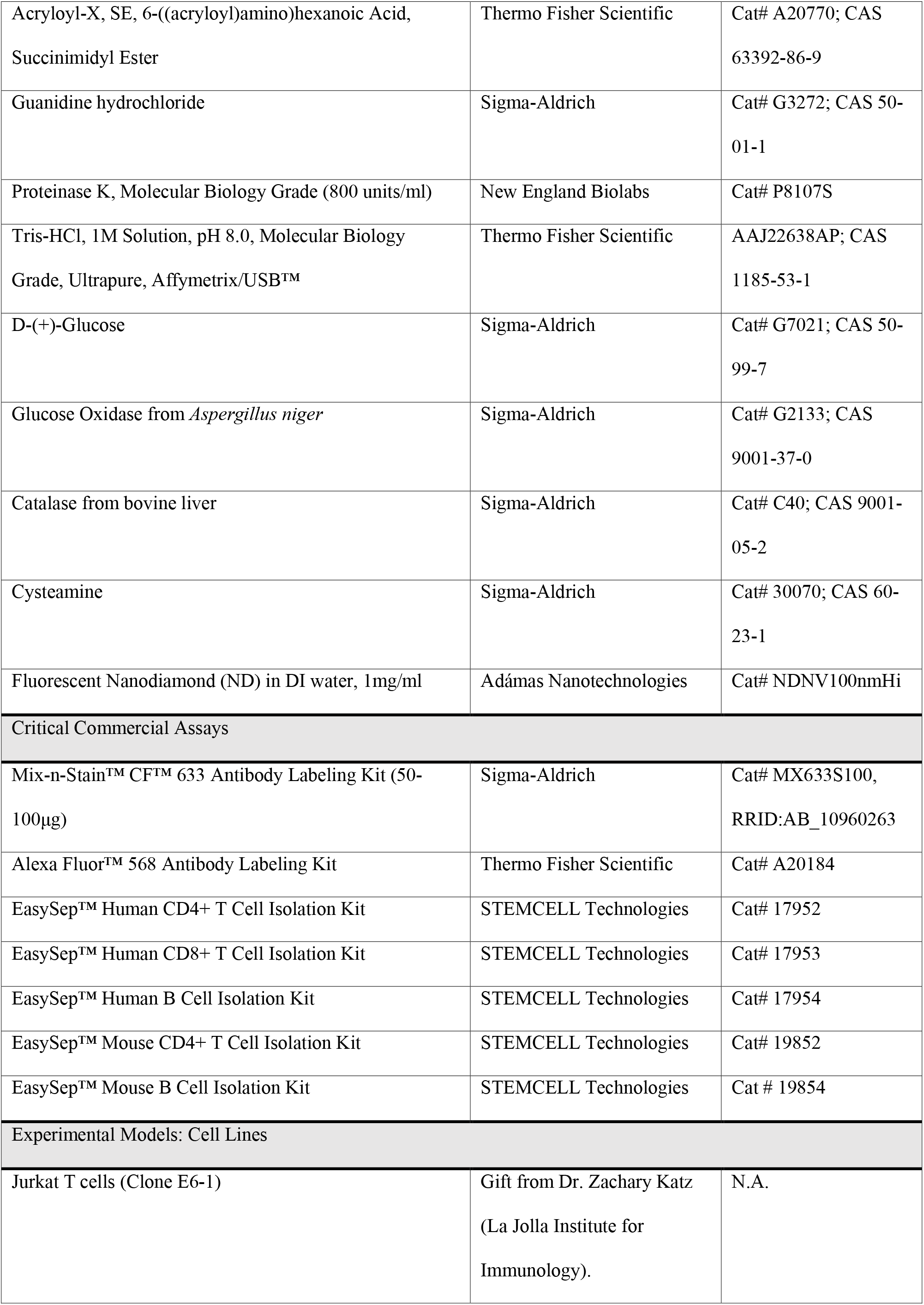
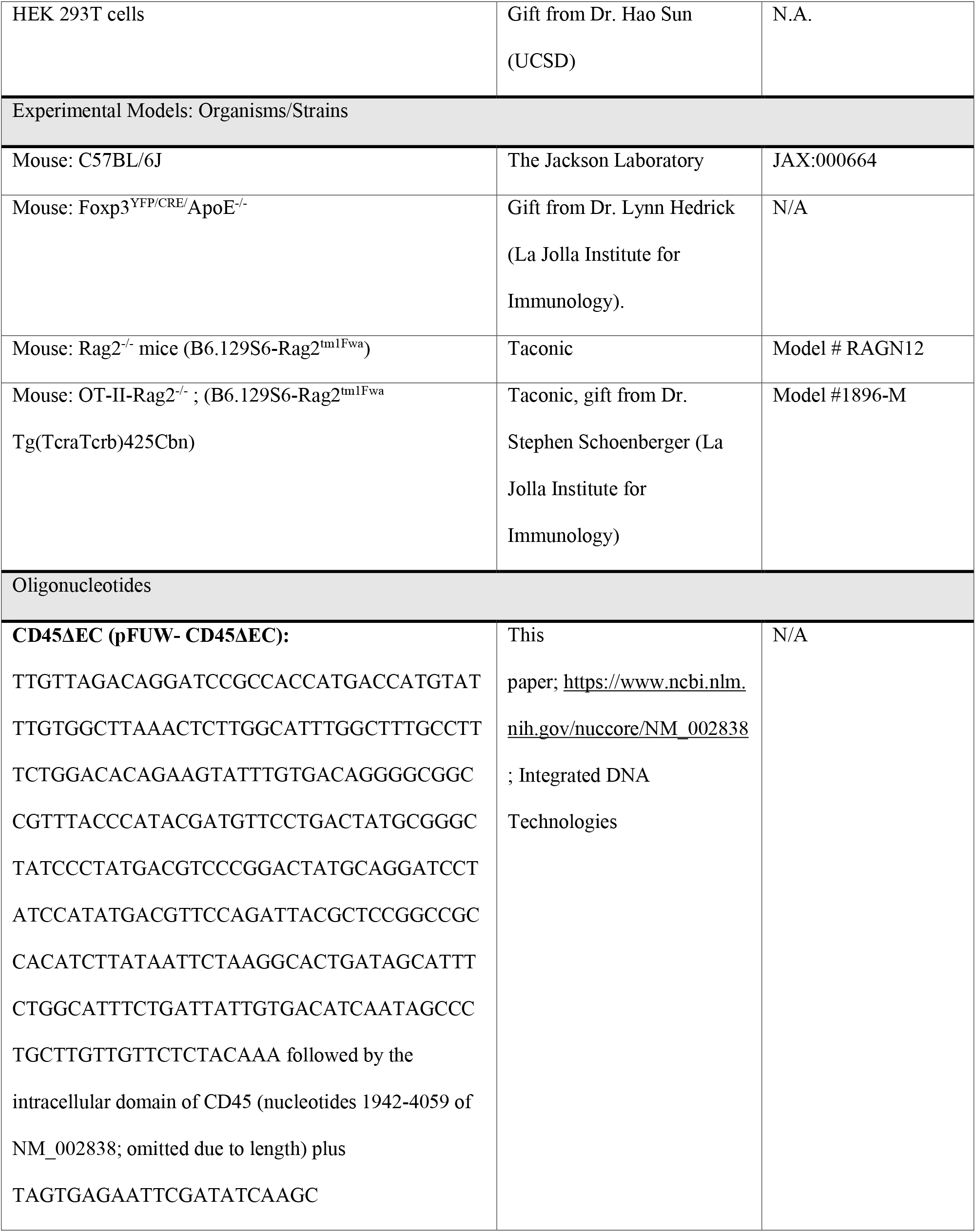
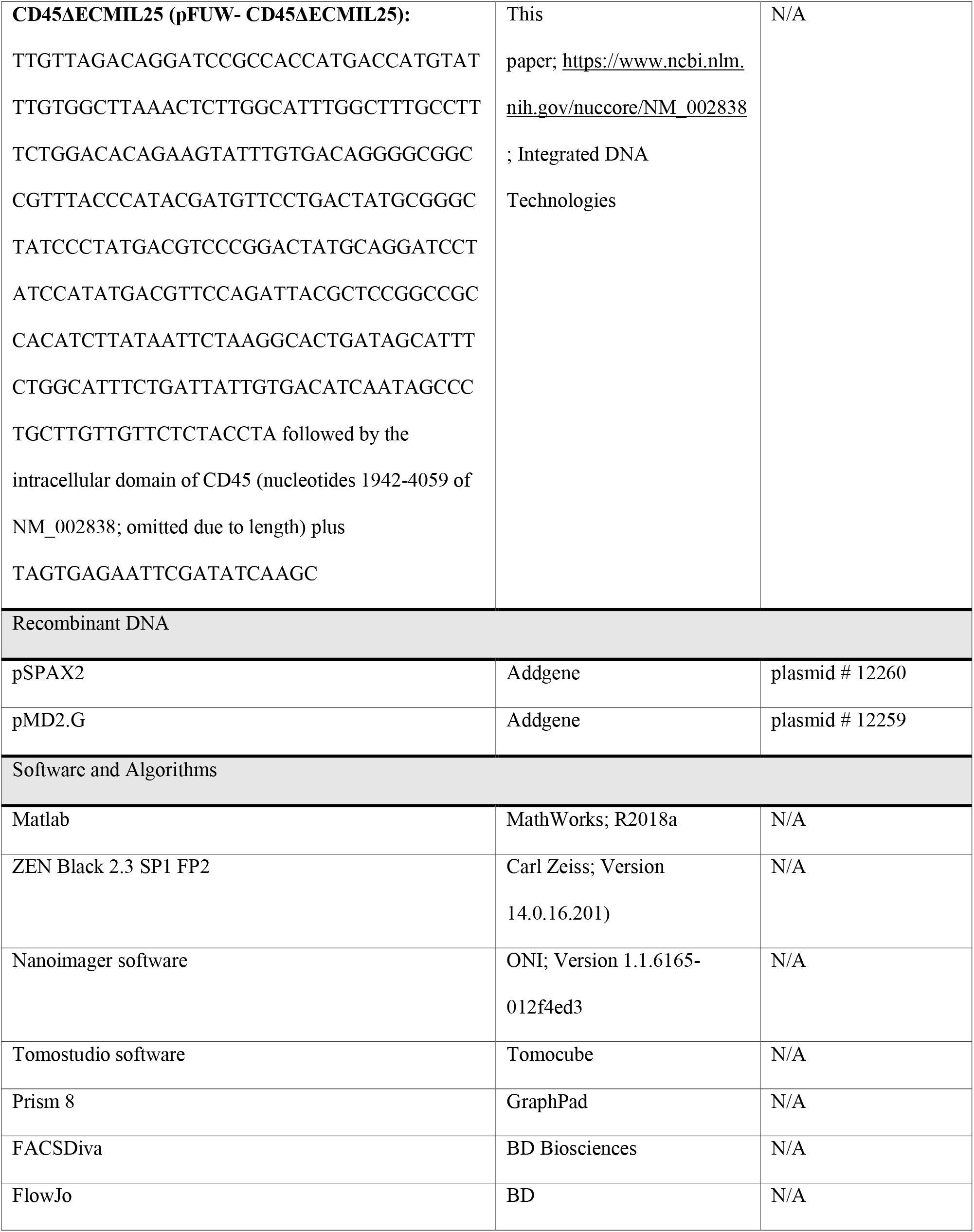

## Supplemental Information

### Supplemental Information titles and legends

**Figure S1.**
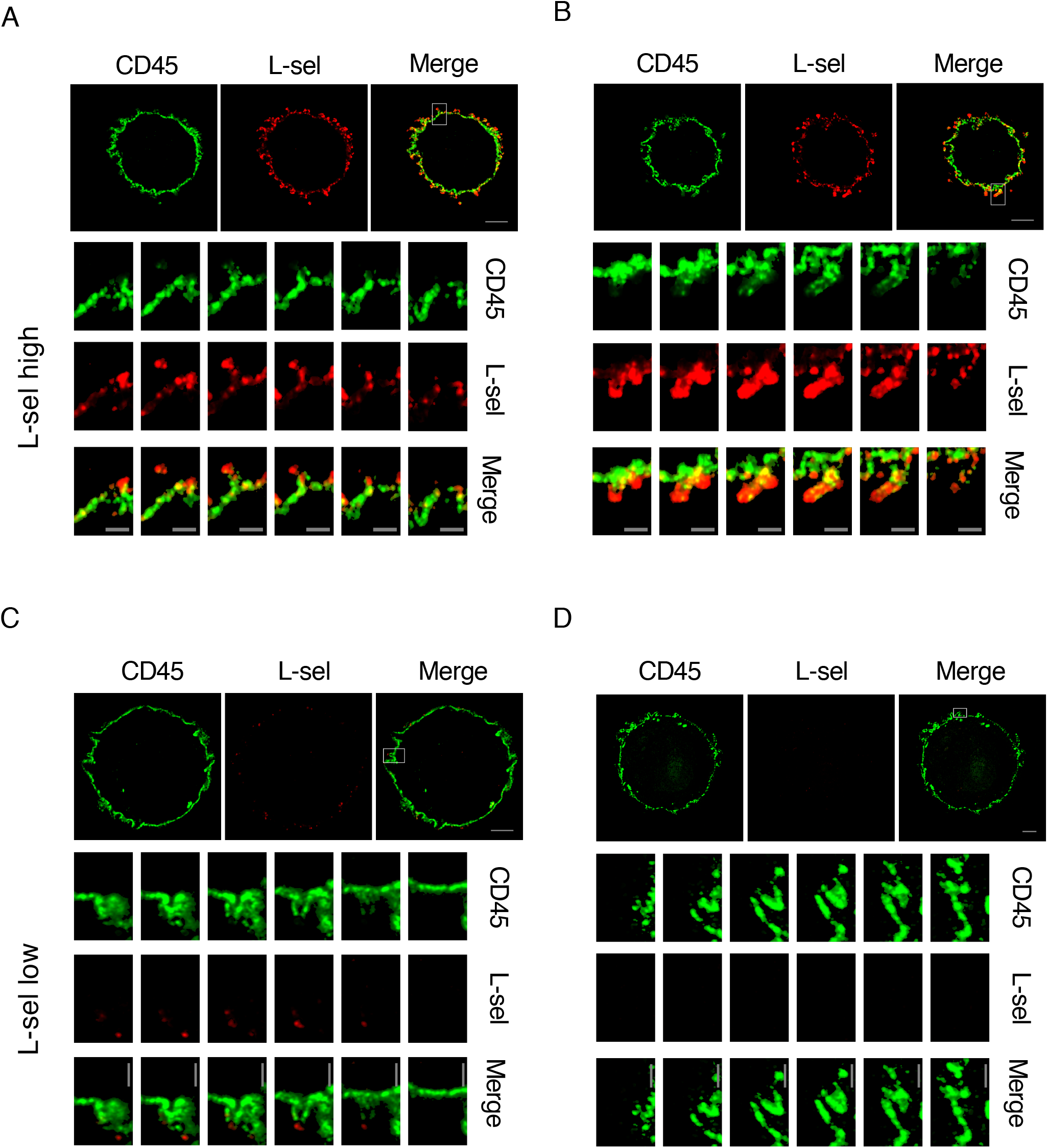
CD45 pre-exclusion in L-selectin-high and -low effector CD4^+^ T cells. Related to Figure 1. (A, B) Representative 4x-ExM-Airyscan images of CD45 (green), L-selectin (red), and the merged images of these two expressed on L-selectin-high *in vitro*-differentiated effector CD4^+^ T cells (top panels). The magnified z-stack series of images with a step size of 125 nm marked by a square in the merged images are displayed in the bottom panels. (C, D) Analysis similar to (A, B) of L-selectin-low effector CD4^+^ T cells. Scale bars: 2 μm in the merged images; 500 nm in the magnified merged images.

**Figure S2.**
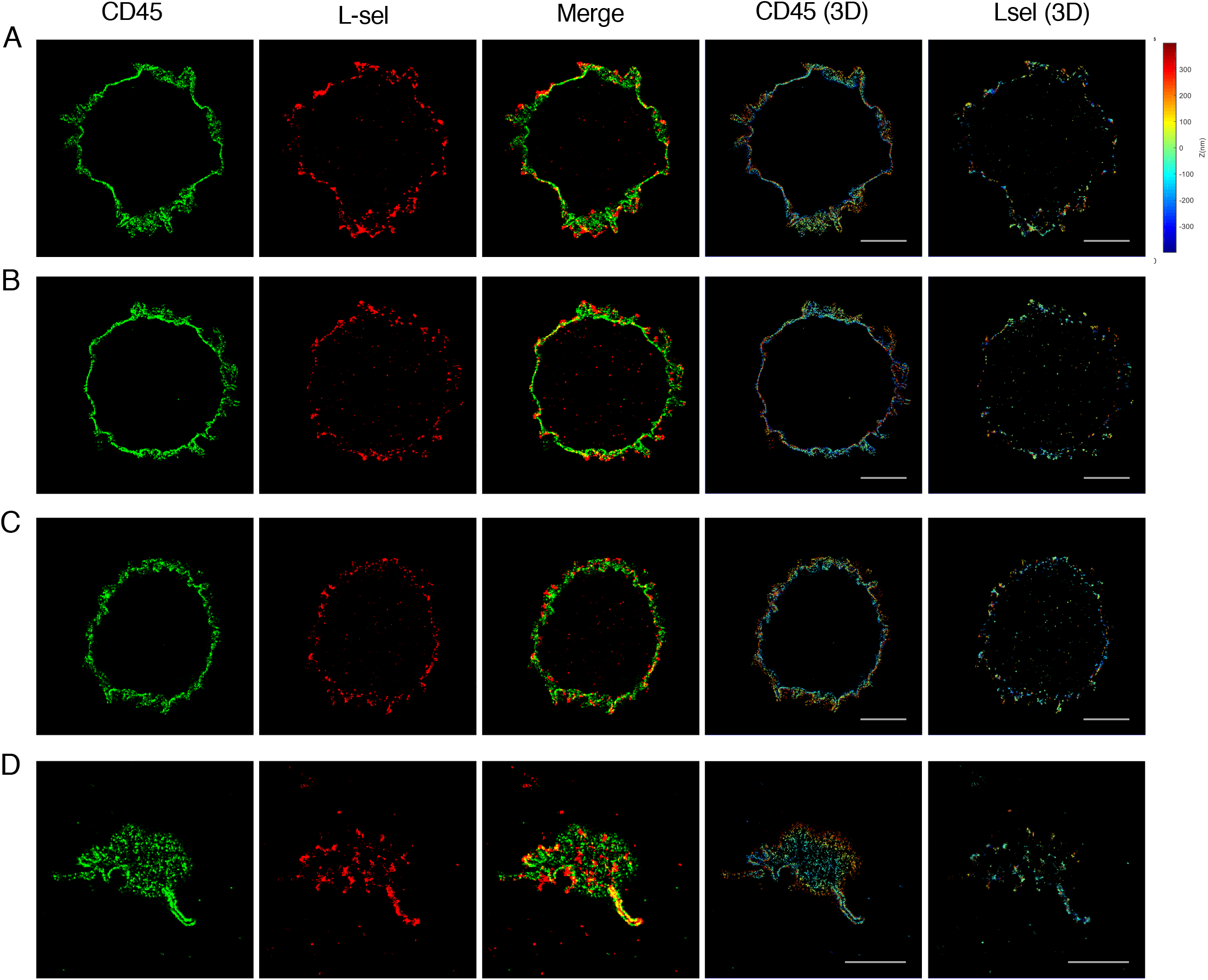
STORM images of CD45 and L-selectin expressed on human CD4^+^ T cells. Related to Figure 1. (A-D) Representative STORM images of CD45 (green) and L-selectin (red) and the merged images of 4 representative cells selected from among 40 cells (3 left rows). The z-dimension localizations of the CD45 and L-selectin molecules were color-coded (2 right rows). Images were acquired at 1.5~2 μm above the glass surface (A-C) or at the glass surface (D). Scale bars: 2 μm.

**Figure S3.**
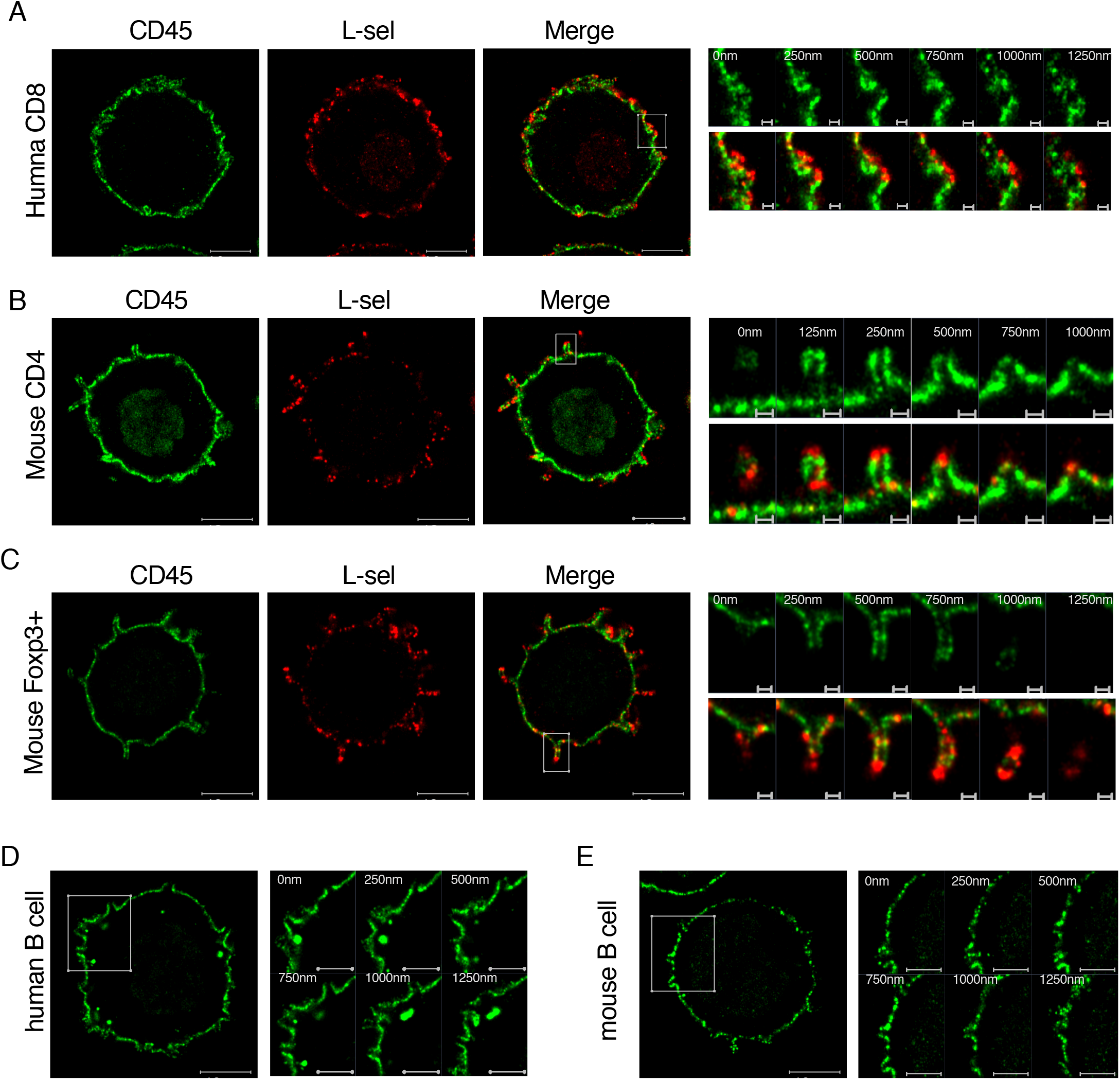
CD45 pre-exclusion from microvilli tips in various types of human or mouse lymphocytes. Related to Figure 1. (A) Representative 4x-ExM-Airyscan image of CD45 (green) and L-selectin (red) and the merged image expressed on a human CD8^+^ T cells. Six z-stack images (step size 250 nm) of the area marked by a square in the merged image were magnified (right). Scale bars: 2.5 μm in left 3 images; 250 nm in the magnified images. (B) Similar analysis of CD45 (green) and L-selectin (red) expressed on mouse CD4^+^ T cells. Scale bars as in (A). (C) Similar analysis of CD45 (green) and L-selectin (red) expressed on a mouse CD4^+^Foxp3-YFP^+^ Treg cells. Scale bars as in (A). (D, E) Similar analysis of CD45 (green) on human (D) and mouse (E) B cells. Scale bars: 2.5 μm in the left images; 1.25 μm in the magnified images.

**Figure S4.**
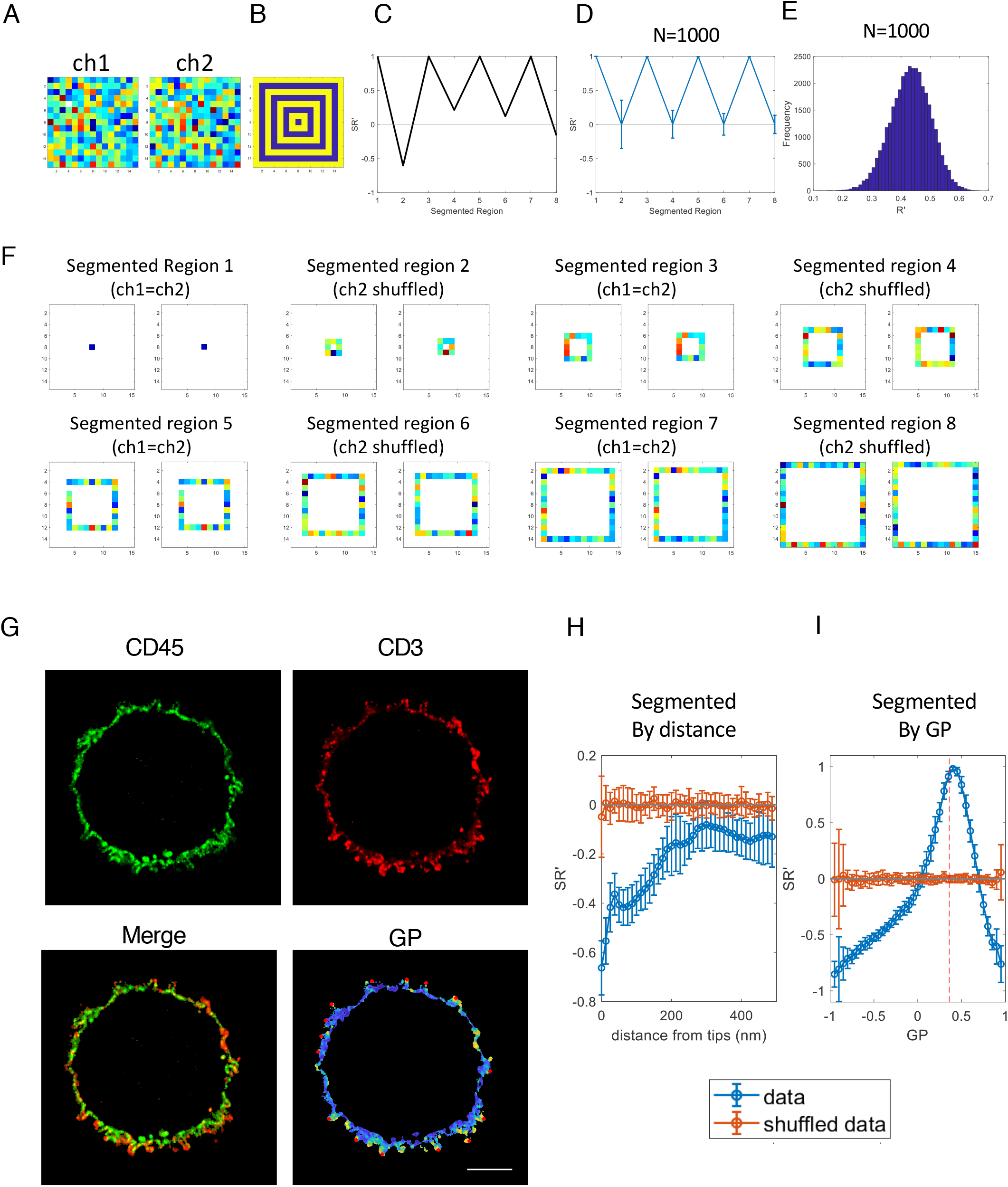
The segment correlation coefficient (*SR*′). Related to Figures 2 and 3; Figures S5-S8; Table S1. *SR’* is calculated between two-channels in the segmented area to present correlation within that area as described in Table 1. (A) An example of simulated images for ch1 and ch2. Ch1 image was generated by assigning random intensities (mean=100; the standard deviation = 2), and ch2 values were identical with ch1 in the blue-marked area in B. The rest of the pixel values of ch1 corresponding to the area marked with yellow in (B) were randomly shuffled and reassigned into the ch2 corresponding to the yellow-marked area in (B). The Pearson’s correlation coefficient, *R’,* between ch1 and ch2 was 0.4. (B) The blue area marks identical pixels between ch1 and ch2 and the yellow area marks non-identical pixels. (C) *SR’* for the eight individual segmented data set of (A) presented in (F). (D) The mean *SR’*s for the eight individual segmented data set similarly presented in (F) from 1,000 simulated data points as (A). The *SR’* within the blue marked area is 1 while the yellow marked area became zero. (E) Distribution of the *R’* between ch1 and ch2 of 1,000 simulated data points. (F) The individual segmented area of (A) calculated as *SR’* for (C). (G) A representative example of a 4x-ExM-Airyscan image of CD45 (green) and CD3 (red) of human resting CD4 T cells and the merged and GP images between the two molecules. The red dots in the GP images represent the location of microvilli tips. Scale bar: 2 μm. (H) *SR’* calculated from 20 frames of the cell presented in (G) segmented by distance from microvilli tips was plotted as a function of the distance (blue) showing that that CD45 and CD3 were negatively correlated near to the microvilli-tips. *SR’* calculated after the two-channel intensities of (G) were randomly shuffled (orange). (I) *SR’* calculated for the data set in (H) segmented by GP values with a step-size of 0.05 were plotted as a function of the GP, showing that the *SR’* became negative where the GPs were near −1 or +1; while the *SR’* became +1 when the GP was close to the mean GP value (0.36±0.92) of the images (red-vertical line). The mean *SR’* was calculated after the two-channel intensities of (G) were randomly shuffled (orange), showing that the mean *SR’* became close to zero. Error bars represent SD.

**Figure S5.**
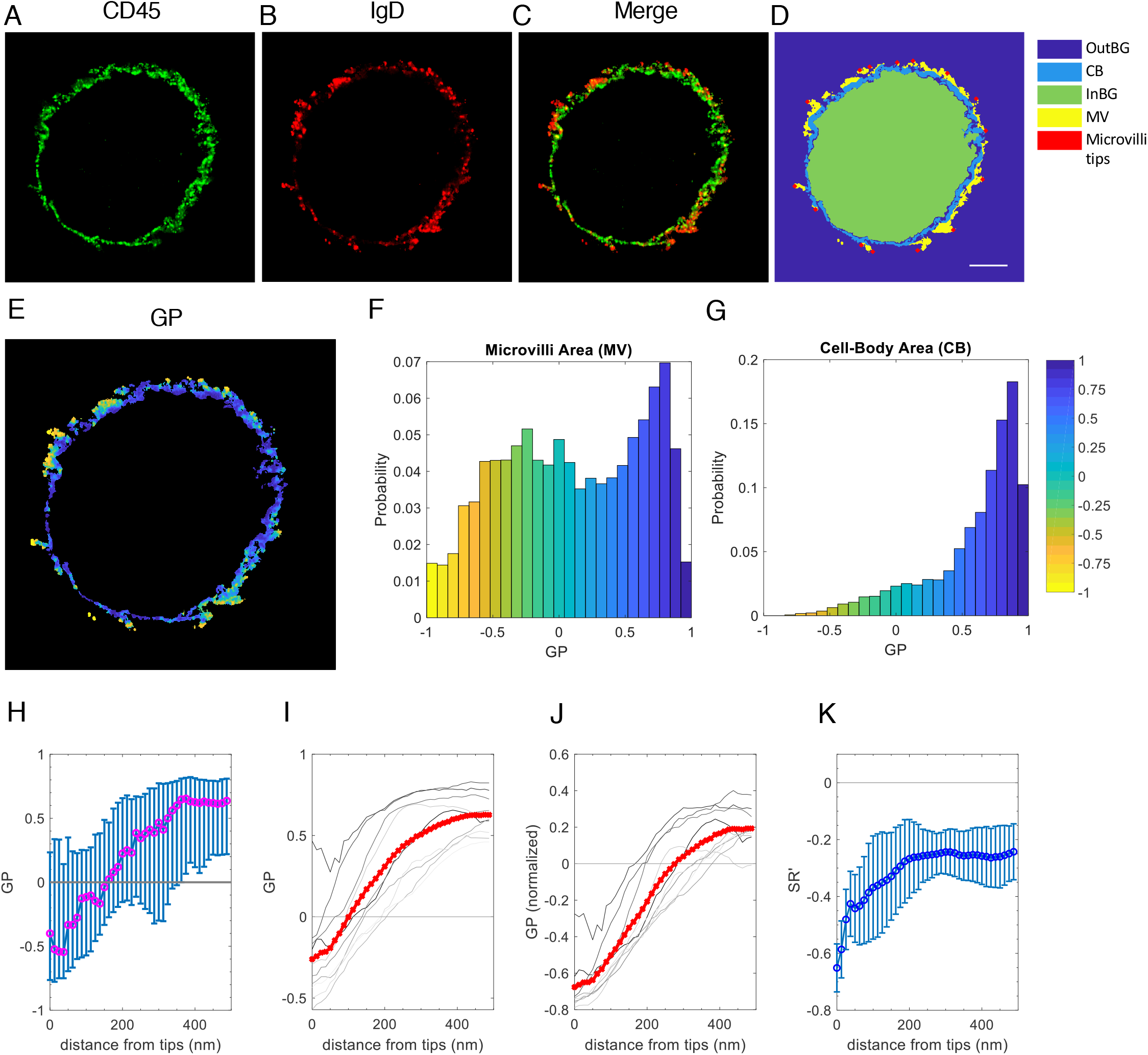
CD45-free membrane-bound IgD molecules on the tips of microvilli of human B cells. Related to Figure 3. (A-C) A representative 4x-ExM-Airyscan image of a B cell labeled with anti-CD45-AF488 (green; A), anti-IgD-CF633 (red; B) and the merged image of the two (C). (D) The area map segmented for OutBG, CB, InBG, MV and microvilli tips of the images in (A-C). Scale bar: 2 μm. (E) The GP image between (A) and (B). (F, G) Normalized distribution of the GP values that correspond to the MV area (F) and CB area (G) presented in (D). (H) The median GP values of image (E) segmented by distance from the tips of microvilli shown in (D) were plotted as a function of the distance from the tips. Error bars represent the 25th percentile and 75th percentile of the data. (I) The median GP values of 10 cells were plotted as a function of the distance from the microvilli tips. Each line represents data collected from 20 z-plane images of a cell; the red line is the mean of the 10 plots. (J) Normalized GP values for each frame were plotted as I. (K) The mean *SR’* values between the CD45 and IgD images segmented by distance obtained from 10 cells were plotted as a function of the distance from the tips. Error bars represent the SD.

**Figure S6.**
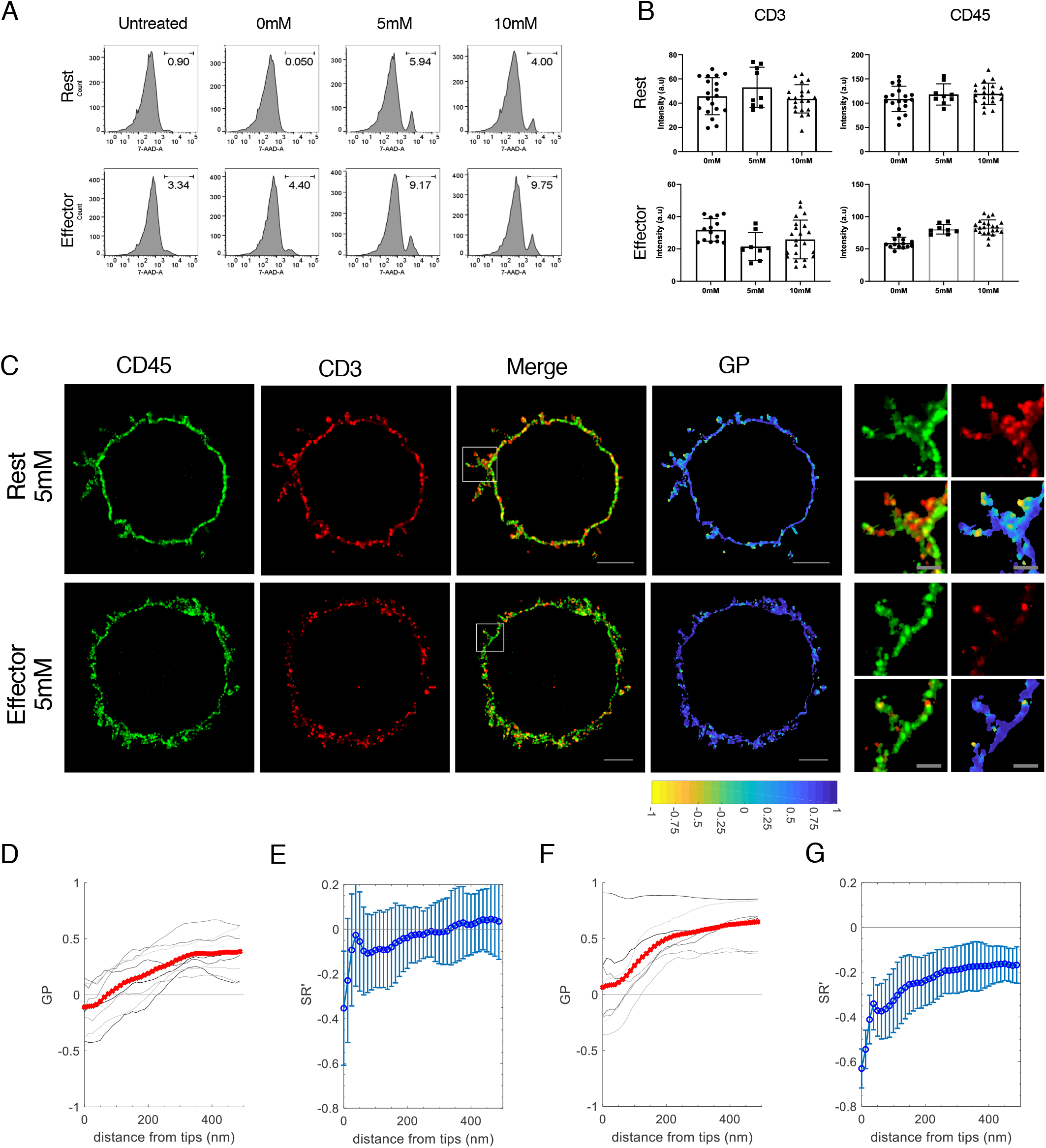
Effect of MβCD on the distribution of CD45 molecules on the surface of CD4^+^ T. Related to Figure 4. (A) The effect of MβCD treatment on the viability of human resting (upper panels) and effector (lower panels) CD4^+^ cells was analyzed by flow cytometry. 7AAD^+^ positive dead cells were counted by FlowJo software. (B) The mean CD45 and CD3 intensities of human resting (upper panels) and effector (lower panels) CD4^+^ T cells treated without (0 mM) or with 5 or 10 mM MβCD. (C) Representative 4x-ExM-Airyscan images of CD45 (green), CD3 (red), and the merged and GP images between the two images of MβCD (5 mM)-treated resting (upper) and effector (lower) human CD4^+^ T cells. Magnified images of the areas marked by a square in the merged images are shown on the right. Scale bars: 2 μm in the merge and the GP images; 500 nm in the magnified images. (D) The median GP values of 9 MβCD (5 mM)- treated resting CD4^+^ T cells were plotted as a function of the distance from the microvilli tips. Each line represents data collected from 20 z-plane images. The red line is the mean of the plots. (E) The mean *SR’* values between the CD45 and CD3 images of the cells in (D) segmented by distance were plotted as a function of the distance from the microvilli tips. Error bars represent SD. (F, G) The median GP values and the mean *SR’* values between the CD45 and CD3 of 8 MβCD (5 mM)-treated effector CD4^+^ T cells were analyzed as in (D) and (E).

**Figure S7.**
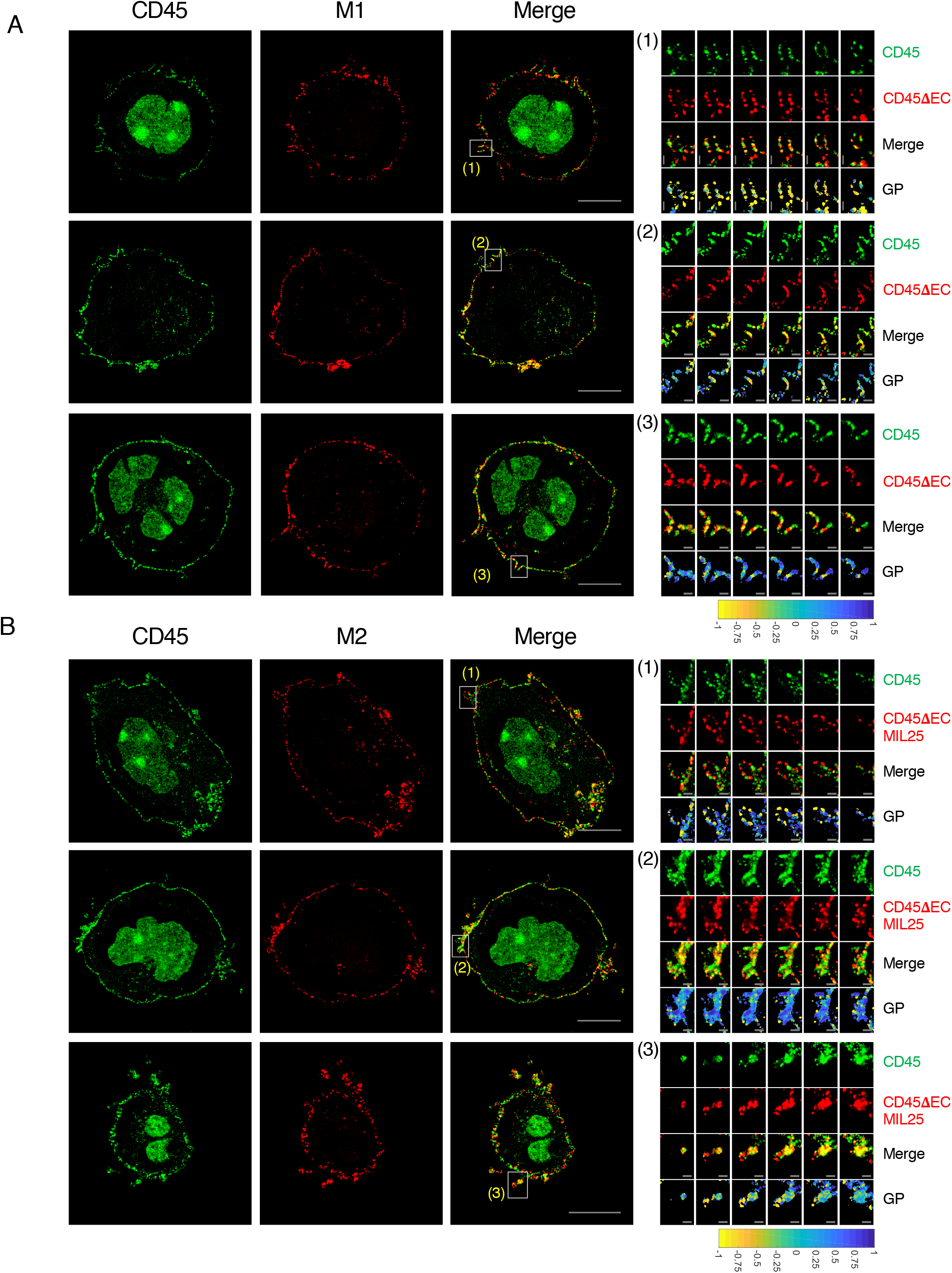
Analysis of Jurkat T cells lentivirally transduced with CD45 mutants. Related to Figure 6. (A) Representative 4x-ExM-Airyscan images of 3 cells out of 15 Jurkat cells lentivirally transduced with CD45ΔEC mutant. The endogenous CD45 images labeled with anti-CD45-AF488 antibody (green), the membrane expressed mutant images labeled with anti-HA-CF633 antibody (red) and the merged images between the two molecules were presented. Scale bars: 5 μm. The six z-stack images (step size 125 nm) of the marked area in the merged images were magnified for each cell (right). Scale bars: 500 nm. The color bar underneath represents GP values between −1 (100% mutant CD45; 0 % endogenous CD45) and +1 (100% endogenous CD45; 0% mutant CD45). (B) Analysis of 3 representative cells out of 25 Jurkat T cells lentivirally transduced with CD45ΔECMIL25 mutant as described as (A).

**Figure S8.**
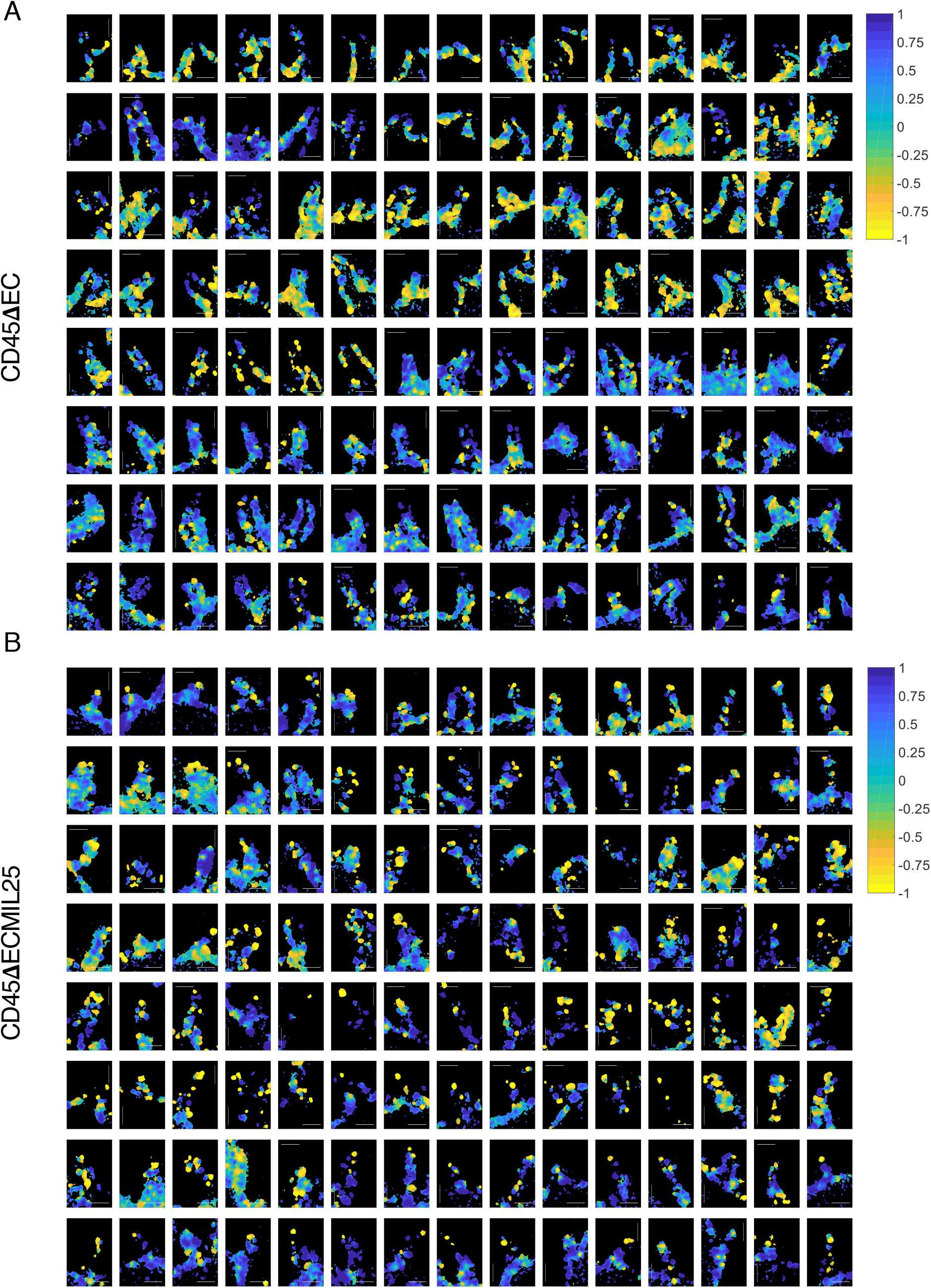
A collection of magnified single microvillus images of CD45ΔEC (A) or CD45ΔECMIL25 (B)-transduced Jurkat T cells. Related to Figures 6 and S7. The color bars on the right represent GP values between −1 (100% mutant; 0% endogenous CD45) and +1 (100% endogenous CD45; 0% mutant). Scale bars: 500 nm.

**Table S1.** Formulas used to calculate GP, *R’*, *SR’*, MOC and MCC. See text for more explanations.

| Abbreviation | Full Name | Equation |  |
| --- | --- | --- | --- |
| GP | Generalized polarization (Yu et al., 1996) | $GP = (X_{ij} - Y_{ij}) / (X_{ij} + Y_{ij})$ | $X_{ij}$ and $Y_{ij}$ are the fluorescence intensity of each pixel indexed $ij$ for channel X and channel Y, respectively. |
| $R'$ | Pearson's correlation coefficient (Rodgers and Nicewander, 1988). | $R' = \frac{\sum_{i=1}^n (X_i - \bar{X})(Y_i - \bar{Y})}{\sqrt{\sum_{i=1}^n (X_i - \bar{X})^2} \sqrt{\sum_{i=1}^n (Y_i - \bar{Y})^2}}$ | n is the total number of data points, $X_i$ and $Y_i$ are the individual data points indexed with $i$, $\bar{X}$ and $\bar{Y}$ are the mean of total X and Y, respectively. |
| $SR'$ | Segment-correlation coefficient | $SR' = \frac{\sum_{j=1}^m (x_j - \bar{X})(y_j - \bar{Y})}{\sqrt{\sum_{j=1}^m (x_j - \bar{X})^2} \sqrt{\sum_{j=1}^m (y_j - \bar{Y})^2}}$ | m is the number of data points within a segmented area. $x_j$ and $y_j$ are the individual data points indexed with j in a set of segmented data, $\bar{X}$ and $\bar{Y}$ are the mean of total X and Y, respectively. |
| MOC | Manders' overlap coefficient (Manders et al., 1993) | $MOC = \frac{\sum_{i=1}^n (x_i)(y_i)}{\sqrt{\sum_{i=1}^n (x_i)^2} \sqrt{\sum_{i=1}^n (y_i)^2}}$ | n is the total number of pixels, where $x_i$ and $y_i$ refer to the intensities of above-threshold pixel value indexed $i$ in the channel X and Y images |
| $MCC_1$, and $MCC_2$, | Manders' correlation coefficient (Manders et al., 1993) | $MCC_1 = \frac{\sum_{i=1}^n (x_i \text{ col})}{\sum_{i=1}^n (x_i)}$<br>$MCC_2 = \frac{\sum_{i=1}^n (y_i \text{ col})}{\sum_{i=1}^n (y_i)}$ | n is the total number of pixels in each image being analyzed, $x_i \text{ col}$ is the pixel intensity of channel X indexed $i$ where the corresponding pixel intensity of channel Y is above the threshold, $y_i \text{ col}$ is the pixel intensity of channel Y indexed $i$ where the corresponding pixel intensity of channel X is above the threshold |

**Table S2.**
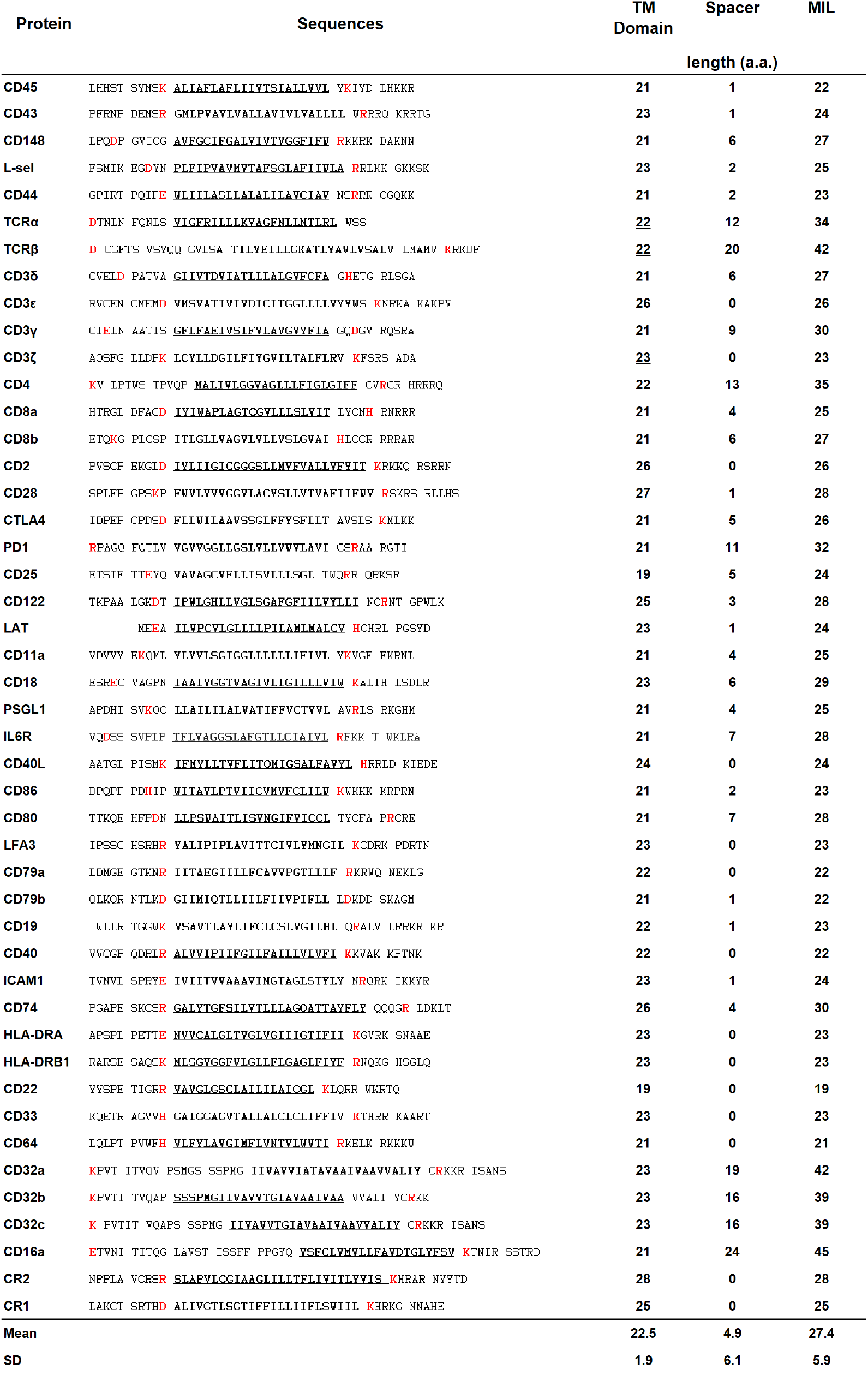
TM domains and the MILs of major cell surface molecules expressed on T cells, B cells or APCs. Related to Figure 5. Amino acid sequences of the TM domain plus flanking residues are shown. The sequences of TM domains predicted in the Universal Protein Resource (Uniport; https://www.uniprot.org/) or reported in references (Call et al., 2006; Krshnan et al., 2016)) are underlined. The nearest charged a.a. residues [lysine (K), arginine (R), glutamic acid (E), aspartic acid (D), histidine (H)] located at each end of the TM domain are shown in red. The length of the TM domain, spacer, and membrane integration limit for each protein are shown on the right. CD45 was excluded from the mean and SD calculation.

**Table S3.**
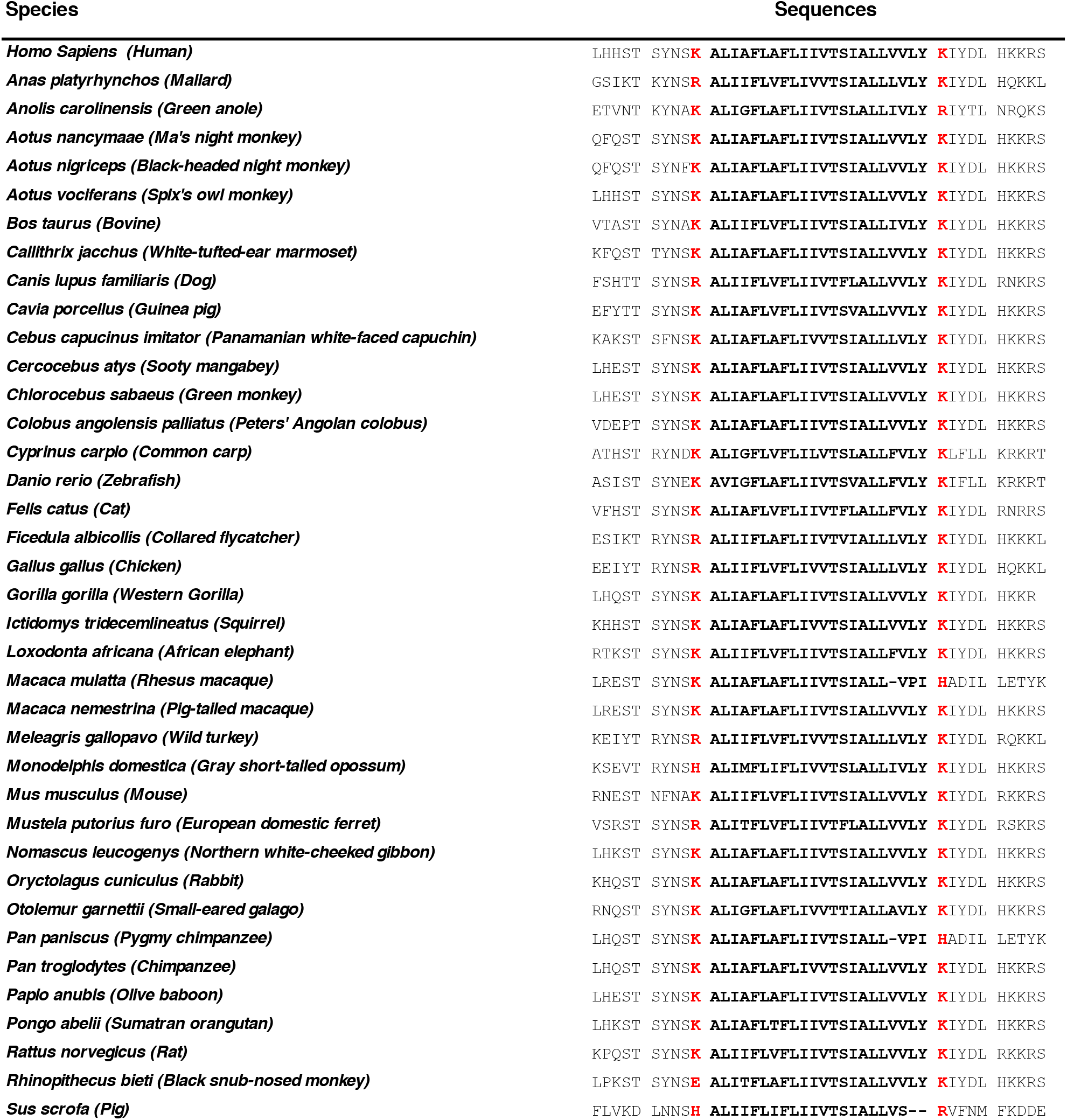
Homologies of the TM domains of 38 CD45 proteins. Related to Figure 5. The amino acid sequences of the TM domains ± flanking residues are shown and the predicted TM domains. Charged TM domain-flanking residues are shown in red.

**Video S1**. Microvilli-mediated interactions between a T cell and an antigen pulsed B cell induce Ca^2+^ influx. (A) A time-lapse of ODT images overlaid with the corresponding Fluo-4 AM images. (B) Fluo-4 AM images of (A). (C) A region of interest (ROI, T cell) of (A) and (B). (D) Mean intensities of ROI area in (B).

**Video S2**. Microvilli-mediated interactions between T cells and antigen pulsed B cells induce Ca^2+^ influx. (A) A time-lapse of ODT images overlaid with the corresponding Fluo-4 AM images (B) Fluo-4 AM images of (A). (C) Three ROIs for T cells in (A) and (B). (D) Mean intensities of ROIs for each cell in (B).

